# SynBot: An open-source image analysis software for automated quantification of synapses

**DOI:** 10.1101/2023.06.26.546578

**Authors:** Justin T. Savage, Juan Ramirez, W. Christopher Risher, Yizhi Wang, Dolores Irala, Cagla Eroglu

## Abstract

The formation of precise numbers of neuronal connections, known as synapses, is crucial for brain function. Therefore, synaptogenesis mechanisms have been one of the main focuses of neuroscience. Immunohistochemistry is a common tool for visualizing synapses. Thus, quantifying the numbers of synapses from light microscopy images enables screening the impacts of experimental manipulations on synapse development. Despite its utility, this approach is paired with low throughput analysis methods that are challenging to learn and results are variable between experimenters, especially when analyzing noisy images of brain tissue. We developed an open-source ImageJ-based software, SynBot, to address these technical bottlenecks by automating the analysis. SynBot incorporates the advanced algorithms ilastik and SynQuant for accurate thresholding for synaptic puncta identification, and the code can easily be modified by users. The use of this software will allow for rapid and reproducible screening of synaptic phenotypes in healthy and diseased nervous systems.

**Motivation:** Light microscopy imaging of pre- and post-synaptic proteins from neurons in tissue or *in vitro* allows for the effective identification of synaptic structures. Previous methods for quantitative analysis of these images were time-consuming, required extensive user training, and the source code could not be easily modified. Here, we describe SynBot, a new open-source tool that automates the synapse quantification process, decreases the requirement for user training, and allows for easy modifications to the code.

## Introduction

Proper brain function relies on the correct wiring of neuronal networks^1–4^. Asymmetric neuron-neuron connections, known as synapses, are the fundamental cell biological units of neural circuits^5–7^. Importantly, synapse loss or dysfunction is a hallmark of many neurodevelopmental and neurodegenerative disorders^8–10^. Therefore, studying synapse connectivity is crucial to understanding how neuronal circuits are established during development, remodeled throughout life, and impacted by diseases.

Synapses are composed of two neuronal structures: the presynapse, located in the axon terminal, and the postsynapse, located on the dendrite. The presynapse contains specialized, neurotransmitter-filled, synaptic vesicles^6,11^. Some of these vesicles are docked at the presynaptic active zone and are fused with the membrane when an action potential reaches the synapse^11^. Neurotransmitters then diffuse into the extracellular space between the pre- and post-synapse, called the synaptic cleft^12^. Neurotransmitters in the synaptic cleft bind post-synaptic neurotransmitter receptors^13–15^. These receptors are transmembrane proteins that pass ions and/or recruit intracellular signaling partners to transduce a signal into the post-synaptic cell^7,13–15^ and are anchored at the postsynapse through a scaffold of postsynaptic density proteins^16,17^.

Synapses are dynamic structures that can be strengthened or lost due to changes in the inputs that neurons receive^18–20^. On the other hand, perturbations in genes that control synaptogenesis would also impact the number and organization of synaptic structures^21–23^. Non-neuronal cell types, such as astrocytes, microglia, and oligodendrocytes, also serve as critical regulators of synapse formation and elimination^24–28^. In particular, astrocytes, the major perisynaptic cells in the brain, strongly induce the formation of excitatory and inhibitory synapses through direct contact with neuronal processes or via the secretion of several synaptogenic proteins (reviewed in ^1,29–31)^.

The gold standard methods for interrogating the structure and function of synapses are electron microscopy and electrophysiology, respectively. Electron microscopy (EM) allows the experimenter to visualize the synapse with a high enough resolution (∼2nm) to resolve the pre- and post-synaptic compartments individually^32–34^. The characteristics of presynaptic vesicles and postsynaptic densities are used to identify excitatory versus inhibitory synapses^34–36^. An experimenter can count the number of opposing pre- and post-synaptic sites in electron micrographs of samples from various conditions to determine if these conditions alter the number of synaptic structures^34–36^.

A common method for investigating synaptic function is the whole-cell patch-clamp analysis of miniature postsynaptic currents. The frequency of miniature postsynaptic currents provides information about the number of synapses a cell receives or the probability of release at the presynaptic site. On the other hand, the amplitude of these currents measures postsynaptic strength^37^.

Electron microscopy and electrophysiology continue to provide high-resolution structural and functional information about synaptic connectivity, but these techniques have major limitations. First, they require extensive sample preparation and specialized equipment, making them difficult to establish in a laboratory that does not specialize in them. Second, they have very low throughput and sample only a small subset of synapses or neurons, making them unsuited for screening multiple experimental conditions.

To address these limitations, higher-throughput methods for assessing synapse numbers using immunostaining have proven useful^38,39^. These histological methods use antibodies to label pairs of pre- and post-synaptic proteins that are specialized to distinct synaptic subtypes. For example, presynaptic markers such as Vesicular Glutamate transporter 1 (VGlut1), Vesicular Glutamate transporter 2 (VGlut2), or bassoon can be paired with postsynaptic markers postsynaptic density protein 95 (PSD95) or Homer-1 to label excitatory synapses^40–44^. On the other hand, the Vesicular GABA transporter (VGAT), together with the postsynaptic Gephyrin, mark inhibitory synapses^41,45,46^. See Verstraelen et al., 2020 for a detailed comparison of synaptic marker performance^47^. When these markers are imaged using light microscopy, due to the resolution limit (200-300nm)^48^, and the short (20-30nm)^44,49^ distance between the pre- and post-synaptic compartments, the color signals appear to overlap at synapses.

Therefore, synaptic structures can be quantified by counting the number of colocalized pre- and post-synaptic markers using specialized image analysis software. One of the first programs for performing this analysis is the Puncta Analyzer plugin for ImageJ^38^, which has been widely used in neuroscience and yielded results that were then validated through EM and electrophysiology^41,45,50,51^.

A major limitation of Puncta Analyzer is that all analysis steps require user input, which are time consuming and can be highly subjective. Thus, image analysis using Puncta Analyzer is a lengthy process that requires extensive user training. In addition, the source code for Puncta Analyzer is complex and difficult to edit to allow for customization.

To circumvent these technical challenges, several alternative analysis pipelines have been developed that automate portions of synapse counting. These include methods to count synapses along a neurite (SynD and SynPAnal) ^52,53^, using additional filtering steps to improve synapse detection from manual thresholding (SynapseJ^54^), using automated thresholding algorithms in FIJI (Synapse Counter)^55^, or development of statistical thresholding algorithms (SynQuant)^56^. While these methods provided important features beyond what is available in Puncta Analyzer, none have been as widely used as Puncta Analyzer, which is well suited to quantify densely packed synapses like those seen in the mouse brain tissues. To circumvent the limitations of Puncta Analyzer, here we developed and tested a new open-source ImageJ-based synapse analysis software we call SynBot, which is optimized not only for in vitro but also for high-noise *in vivo* images and allows for a wide variety of options to tailor the analysis to the experimenter’s needs. We compared the accuracy and efficiency of SynBot with Puncta Analyzer using simulated and experimental data which were previously validated by EM and electrophysiology ^57,58^.

## Results

### Synapse labeling by immunohistochemistry

Synapse quantification relies on the use of immunohistochemistry to fluorescently label the pre- and post-synaptic compartments (**Figure 1A**). A standard immunostaining workflow is used, including 1) fixation and permeabilization of the sample, 2) application of primary antibodies against at least one pre-synaptic and one post-synaptic marker, 3) application of fluorescent secondary antibodies against the species of the primary antibodies used, and 4) image acquisition by fluorescence microscopy (**Figure 1B**). This protocol results in images where either excitatory (**Figure 1C**) or inhibitory (**Figure 1D**) synapses can be visualized. The details of the methods are explained in the Methods section and at protocols.io (https://www.protocols.io/view/synbot-protocols-3byl4qewjvo5/v2).

**Figure 1:**
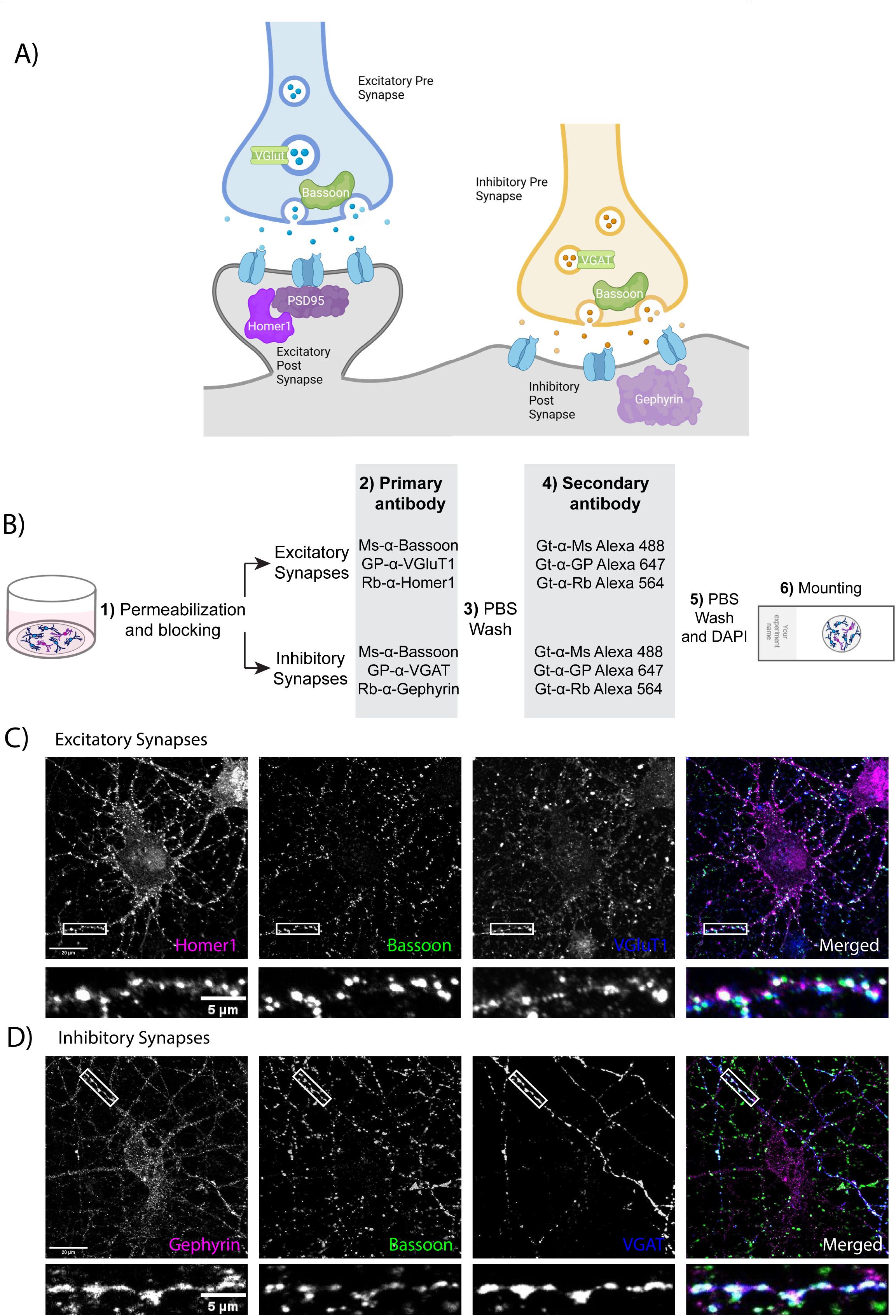
Labeling of synapses as apposing colocalization of pre- and post-synaptic markers. a. Illustration of excitatory and inhibitory synaptic structures. Presynaptic markers used in this paper are shown in green and postsynaptic markers are shown in magenta.
b. Synaptic staining workflow.

1. After fixation with 37°C PFA 4%, the neurons are permeabilized and non-specific epitopes are blocked in antibody buffer containing 50% NGS and 0.2% Triton X-100.
2. Neurons are incubated with primary antibodies targeting pre- and post-synaptic compartments of excitatory or inhibitory synapses.
3. After overnight incubation with primary antibodies, the neurons are washed with PBS.
4. Neurons are incubated with secondary antibodies conjugated to Alexa fluorophores.
5. After 2 hours of room temperature incubation with secondary antibodies, neurons are again washed with PBS and the neuronal nuclei are stained with DAPI.
6. Coverslips are mounted on glass slides using a medium that protects from bleaching of Alexa fluorophores for imaging. For details, check the methods section and protocols.io.
c. Representative image of excitatory synapses made onto a cultured cortical neuron stained with Homer1, Bassoon, and Vglut1. The scale bar in the larger image represents 20 μm, and the scale bar in the smaller image represents 5 μm.
d. Representative image of inhibitory synapses made onto a cultured cortical neuron stained with Gephyrin, Bassoon, and VGAT. The scale bar in the larger image represents 20 μm, and the scale bar in the smaller image represents 5 μm.

### Developing an automated synapse quantification tool

Our first goal in developing SynBot was to enable the efficient quantification of large imaging datasets. SynBot combines many of the image processing and analysis steps that were performed for analysis with Puncta Analyzer into one automated workflow. Toward this aim, we first developed an ImageJ macro to automate the pre-processing steps that convert raw images into numerical representations, which can be used to calculate synaptic colocalizations (**Figure 2**). First, SynBot checks the image files to determine if they are z-stacks of confocal images or single images (either single optical section from a confocal or epifluorescence image). Then SynBot processes the z-stack files to produce max projections of each 1µm stack. This step is of particular use for *in vivo* synapse number analyses, which will be discussed in detail in a later section. Second, all images are converted to the RGB format, the color format Image J uses. Third, based on user selection, the FIJI Subtract Background, and Gaussian Blur Filter functions are applied to remove noise from the image. The fourth step is the thresholding of individual synaptic puncta to set a value for the background of the image and exclude the pixels in the image that have an intensity value less than the background. A threshold value can be determined in several different ways. The most common ones are manual thresholding and ilastik-based automated thresholding. The users can also set the threshold to a specified value. Fifth, the user decides whether to quantify synapses within a region of interest or within the whole image. After entering these user inputs, the program runs the FIJI Analyze Particles function to record the location and area of each punctum in the region of interest of the thresholded image. This function records the X and Y coordinates and area of each punctum for each color channel. Each punctum is then compared to the puncta in the other channel of the image by approximating each punctum to a circle or through a pixel-by-pixel approach discussed below.

**Figure 2:**
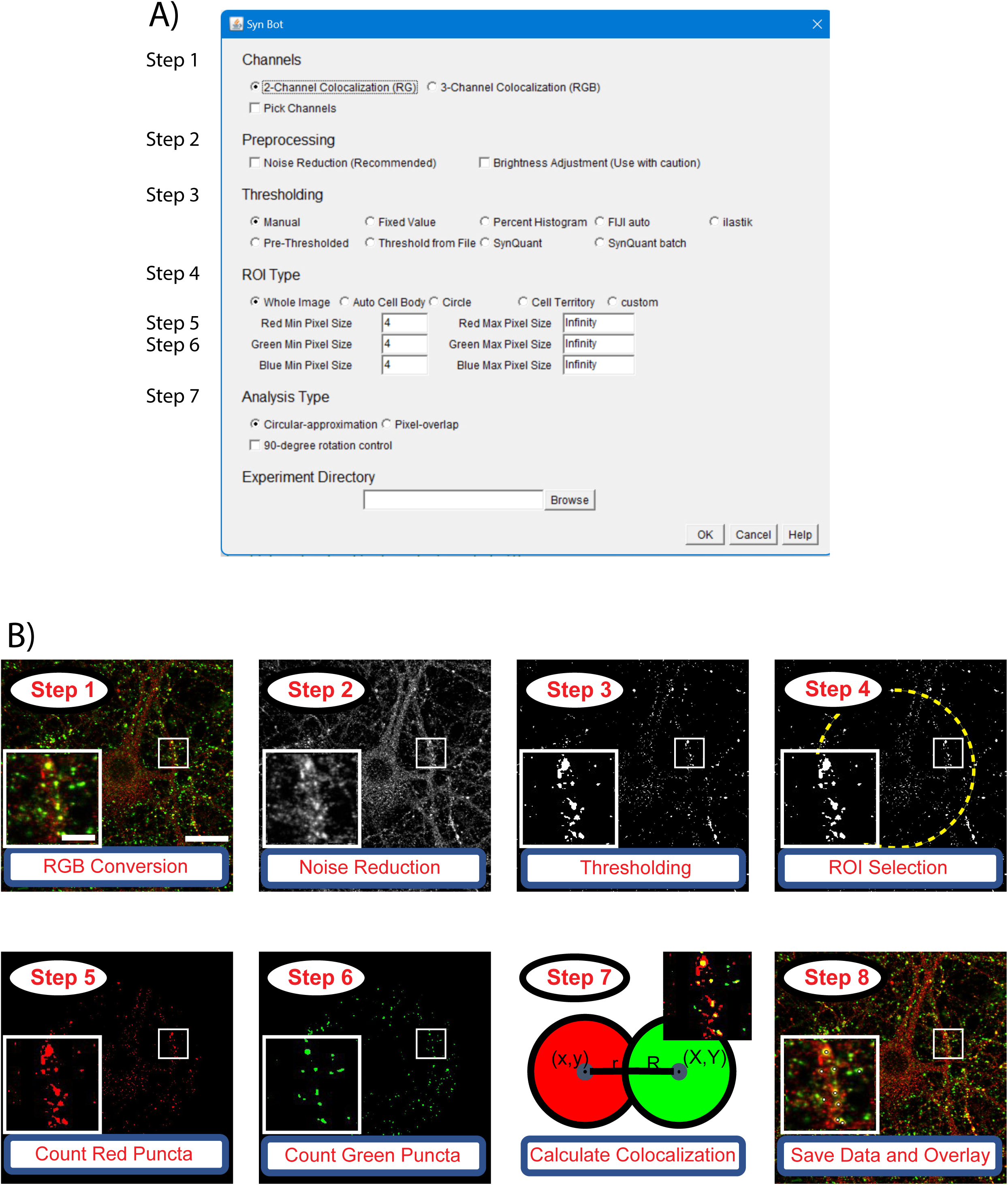
Workflow for SynBot synapse quantification. a. The SynBot selection menu serves as the primary point of user input to adjust options for synapse analysis.
b. The main steps in the SynBot colocalization analysis:

1. Images are loaded into the software and converted to RGB.
2. Images are denoised using the FIJI Subtract Background and Gaussian Blur Filter functions.
3. Each image channel is thresholded to separate synaptic puncta from the background.
4. A region of interest (ROI) for colocalization quantification can be selected (yellow dashed-line).
5. 5-6) Red and green puncta are detected by FIJI’s Analyze Particles function and counted.
6. Colocalization between red and green puncta are calculated. See Figure 3.
7. The colocalized puncta are overlayed on the original image, and the data are saved as a .csv file. The scale bar in the larger image represents 20 μm, and the scale bar in the smaller image represents 5 μm.

### SynBot analysis workflow

The files required for running SynBot and the installation instructions can be found at https://github.com/Eroglu-Lab/Syn_Bot. There are also tutorials and instructional videos on protocols.io (https://www.protocols.io/view/synbot-protocols-3byl4qewjvo5/v2). Once the Syn_Bot.ijm macro is started, a selection menu is displayed to allow the users to select their analysis parameters (**Figure 2**). The selection menu has 6 main sections: 1) Channels, 2) Preprocessing, 3) Thresholding, 4) ROI type, 5) Analysis type, and 6) Experiment Directory. The detailed functions of each option on the SynBot selection menu are defined in **Table S1**. Below, we summarize the utility of each section.

**1) Channels:** The fundamental objective of SynBot is to determine the overlap between different color channel objects (i.e., pre- or post-synaptic puncta). SynBot does this by first converting images into the red, green, and blue (RGB) format, which allows for consistent labeling of channels throughout the program. Users can select either the 2-channel option to analyze colocalization between the red and green channels or the 3-channel option to analyze colocalization between the red, green, and blue channels. If your images are in CMYK (cyan, magenta, yellow, and key (black)) format, or you would like to analyze red and blue or green and blue puncta colocalization, then there is also the “Pick channels” option. Z-stack images are max projected to produce individual projections that SynBot analyzes as separate images. For the experiments described below, every 3 Z-stacks were combined to produce a 1 um max projection. The number of stacks to project together can be adjusted by the user or set to 1 to allow individual processing of each optical section.
**2) Preprocessing:** Images can be preprocessed after RGB conversion to reduce noise or adjust the brightness. Noise reduction is performed by applying FIJI’s subtract background plugin followed by a Gaussian blur filter to aid in object detection. This process removes some of the image background to aid in object detection (see https://imagej.net/plugins/rolling-ball-background-subtraction). Brightness adjustment changes the intensity values of each image such that there is an equal percentage of saturated pixels across images. As these methods modify the image data that SynBot will use for analysis, we advise users to apply them with caution and ensure they are producing accurate results. These modifications should also be consistent across groups of images that will be included in the same experiment.
**3) Thresholding:** One of the most challenging aspects of image analysis is thresholding, the process of distinguishing the foreground (i.e., synaptic puncta) of an image from the background. To discriminate between true synaptic puncta and the background noise is not trivial due to the small size of synaptic puncta and variability in staining and imaging parameters between users and microscopes. Therefore, here we include a number of possible options that SynBot users can utilize. The most popular of these options is manual thresholding, where the user selects a threshold value for each channel of each image, and SynBot runs the rest of the analysis. In that sense, this option is similar to Puncta Analyzer, the predecessor of SynBot. However, we designed Synbot to have several time-saving user-friendly features. For example, Synbot automatically displays each of the image channels to be thresholded, minimizing the time required for user input. However, the manual thresholding method requires extensive user training to use properly and necessitates a significant amount of hands-on user time. To address the limitations of manual thresholding, we implemented several automated thresholding methods. First, we created a new ilastik-based^59^ thresholding method, where a machine learning model is used to threshold each channel of each image. ilastik is an open-source program developed by Anna Kreshuk’s lab at the European Molecular Biology Laboratory. ilastik is trained by the user in a small number of images (typically 3-5) to threshold by extracting a set of user-defined features from the image and then feeding these features into a random forest machine learning model. This model is then applied to the rest of the images in the data set to determine if a given pixel is part of the foreground or background of the image. As in all thresholding methods, ilastik requires careful troubleshooting to ensure accurate results. The training should be done for each experiment independently and should not be used across datasets unless they are collected under identical conditions, such as replicates of the same experiment performed by the same investigator. However, using ilastik has a distinct advantage: it can be applied to a large number of images without the need for further user input and only takes 5-15 minutes to generate the trained model. In addition to ilastik, we also integrated the SynQuant algorithm for thresholding into SynBot. This method was developed by Wang et al. in 2020^56^, to specifically threshold synapses from immunofluorescence images through a statistical probability-based approach. This is explained in detail in Wang et al., 2020 but in short, the method assesses each potential synaptic object by comparing it to its neighboring pixels based on fluorescence intensity, object size, and local contrast. The SynQuant implementation within SynBot allows the user to adjust the following parameters: Z-score threshold (a statistical measurement of how many standard deviations values are from the mean of a population), minimum object size, maximum object size, minimum object fill, maximum width to height ratio, and estimated noise standard deviation. For most datasets, the noise standard deviation and Z-score threshold are the only parameters that require adjustment. The SynQuant batch processing of images through SynBot takes approximately 5-10 minutes for the size of data sets presented here without any need for user input, so trying different parameters on the same images is feasible and recommended to determine the most accurate conditions for analyses. We added another feature to Synbot to aid the reproducibility of results between users. All the thresholding values are saved for each image analyzed. Therefore, an analysis performed with SynBot can be efficiently reproduced later using the “Threshold From File” option. This option asks the user to supply a CSV file containing the threshold values for each channel of each image. This option does not work for the ilastik or SynQuant thresholding options because these algorithms do not use a set thresholding value, but they can already be easily reproduced since they are unsupervised methods.
**4) ROI Type:** SynBot’s colocalization analysis can be targeted to a region of interest (ROI) within the input image. One common application is the restriction of the analysis to the region around the soma of a cultured neuron, as in Figure 5. Alternatively, users can implement their own complex ROIs by using the FIJI clear functions to remove unwanted regions prior to running SynBot on the entire image. During the selection of the ROI, the user is also asked to input a minimum and a maximum pixel size for each channel puncta. The area of the ROI selected for each image is recorded and included in the summary.csv output file.
**5) Analysis Type:** Puncta analyzer calculated colocalization based on a circular approximation colocalization approach^38^ (**Figure 3**). Therefore, we included this option also in SynBot. With the circular-approximation analysis mode selected, the coordinates and radius of each punctum for the channels to be analyzed are passed through the Syn_Bot_Helper java plugin (packaged within https://github.com/Eroglu-Lab/Syn_Bot). We used a Java plugin because it can perform the necessary calculations much more efficiently than if they were done with ImageJ macro language. These values for each punctum in the first channel are then compared to each punctum in the second channel to detect colocalizations and calculate their area using the geometry summarized in **Figure 3**. First, the distance between the centers of the two circles is calculated (**Figure 3A**). If this distance is less than the sum of the radii of the two circles, then they are counted as a colocalization, and the area of the colocalization is calculated (**Figure 3B**). The basic idea of the area calculation is to use the two intersection points of the circles along with the center of each circle to define two triangles while also finding the areas of the sector of each circle between the two intersection points. Subtracting the area of each triangle from the area of its surrounding sector gives half of the overlapping area contributed by that punctum. This can be done for both puncta to give the total area of the overlap (co-localization)^60^. The coordinates and area of each colocalization are then stored and added to the colocalized puncta count. This method would only work if the synaptic puncta had high circularity, which can be determined using the Measure tool in FIJI and is recorded for individual puncta by SynBot. In our experience, most synaptic markers used in primary culture, or the mouse cortex are approximately circular. A table of experimental circularity values for the antibodies used in this paper is provided in Table S2. We also directly compared this circular approximation to the pixel-by-pixel alternative and found nearly identical results for the types of images shown here (Figure S3). The circular-approximation method is too slow to practically analyze three channel images and fails to correctly calculate colocalizations between non-circular objects. To overcome these limitations, here we developed a pixel-overlap colocalization method within SynBot (**Figure 3C**). Rather than approximating each punctum to a circle, this analysis mode compares puncta in a pixel-by-pixel fashion using the FIJI Image Calculator plugin’s AND function. The AND function generates a new image that contains pixels that overlap between the red punctum and a green punctum. This new image can then be used to quantify co-localization with a third channel (blue). The coordinate and area information for colocalized puncta is then collected. Thus, this method is more accurate and versatile and runs more quickly than the circular approximation method.
**6) Experiment Directory:** The final user input field for the SynBot selection menu is to pick a folder that contains experimental images, hereafter called the experimental directory. The directory folder should include at least one subfolder. The subfolder(s) of the experimental directory should contain only the image files to be used by SynBot. We often divide experimental groups between the subfolders of the experimental directory, but this is not required and does not affect the analysis. It is also important that image names contain only one “.” character preceding their file extension. The program uses this “.” to separate the file extension from the image name and will not run properly if there are multiple. Incorrect file naming can be especially problematic with some image type conversion programs that name each image with both the old and new extension (e.g., converting a Leica .lif file to .tif format can result in a name like “image1.lif.tif”).

**Figure 3:**
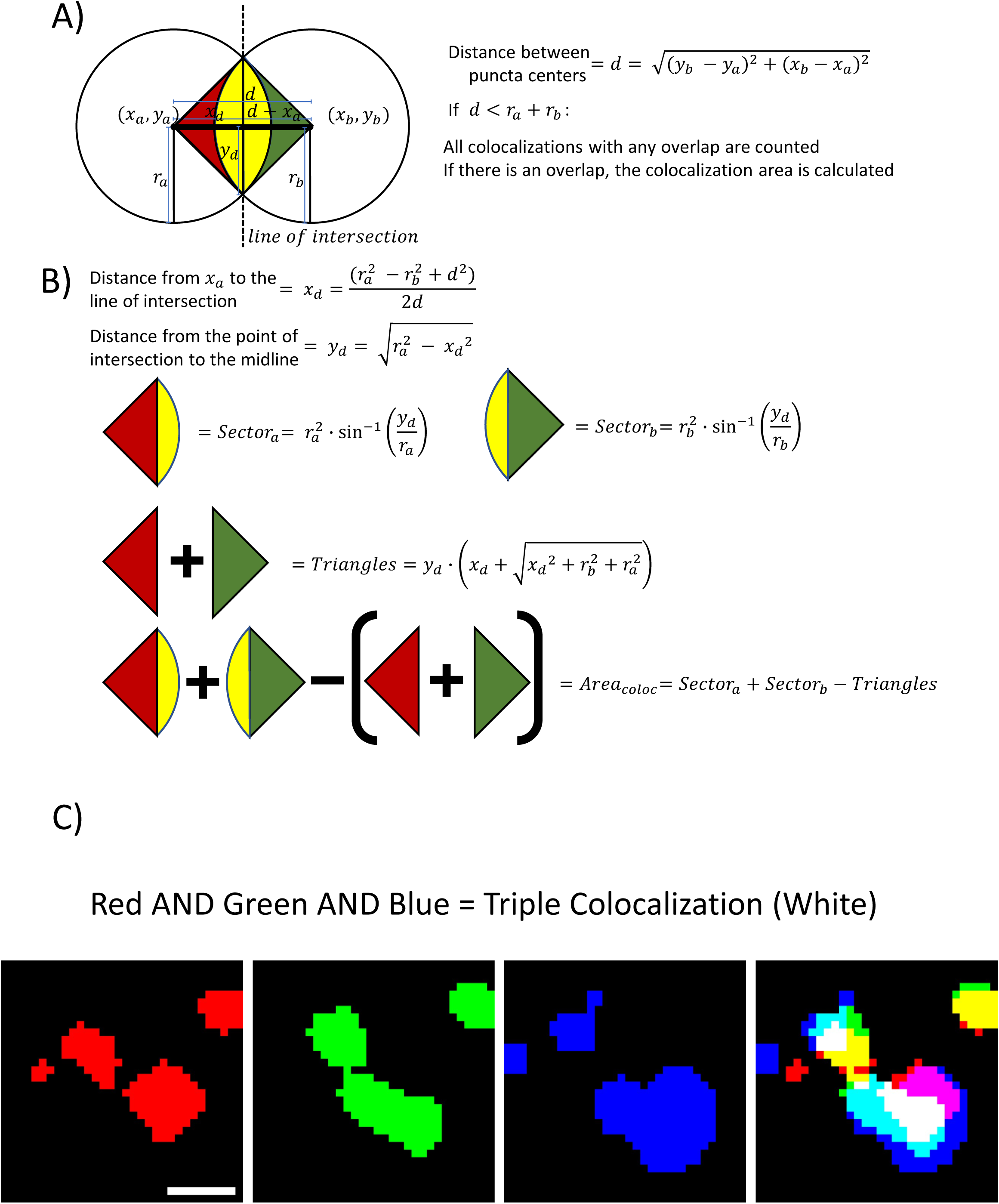
Detection and calculation of puncta colocalization. a. SynBot detects colocalizations between synaptic puncta by first calculating the distance between the centers of the puncta and then determining if that distance is less than the sum of the radii of the puncta. If this distance is less than the sum of the radii, the colocalization is recorded and the area of colocalization is calculated.
b. SynBot runs the following calculations for the area of colocalization area.

1. Calculate the distance between the center of the first puncta and the line of intersection.
2. Calculate the distances from the midpoint of the distance between the two puncta centers (d) to the intersection of the edges of the two puncta.
3. Calculates the area of the sector of the first puncta between the two intersection points.
4. Calculates the area of the sector for the second puncta.
5. Calculates the area of the rectangle formed by the two centers and intersection points; the sum of the two triangular portions of the sectors that are not part of the overlap.
6. The general form for the area of colocalization is the sum of the two sectors minus the two triangular portions outside of the overlap.
c. Pictorial representation of the pixel-overlap analysis mode where FIJI’s image calculator is used to identify pixels that are part of both the red and green channel for a 2-channel colocalization or part of the red, green, and blue channels for a 3-channel colocalization. Scale bar represents 1 μm.

### SynBot Outputs

A key feature of SynBot is that the users can easily check if the method is counting the correct objects as colocalized puncta because after each image is analyzed, SynBot creates and saves a feedback image within the Output folder using the name of the image with “_colocs” appended to the end. This “_colocs” image shows the original input image with white circular overlays at the center position of each recorded colocalization. New SynBot users are strongly encouraged to check these images after running the macro and adjust their thresholding parameters or retrain their ilastik model if they see too few or too many calculated colocalizations that do not match what is evident by eye. The thresholded images for each channel are also saved so the users can later validate whether each channel was accurately thresholded.

The primary output of SynBot, which contains the numbers of colocalized puncta per image, is the “Summary” csv file. This file contains the most commonly used summary measurements for synapse counting, such as the total numbers of red, green (and blue if triple colocalization is used), and the numbers of colocalized puncta from each image. The file also includes the thresholds and the settings for minimal pixel size. To allow for analysis of the cumulative properties of synaptic puncta, the macro also saves the X and Y coordinates and area of each individual puncta for the red, green, blue, and colocalized puncta into separate CSV files. These results are then ready for statistical analyses by using other software such as R or GraphPad.

### Quantification of synaptic colocalization by SynBot in simulated images

To validate SynBot’s performance and compare the automated thresholding methods that are integrated within SynBot, we tested SynBot on simulated synapse images. We produced 20 simulated images with 667 red puncta, 666 green puncta, and 333 points where these puncta overlap to form a synapse. We then added a Gaussian noise background to each image with a mean and standard deviation proportional to the intensity histogram of the original image multiplied by 0.00, 0.25, 0.50, 0.75, or 1.00 to simulate images with varying noise levels (**Figure 4A**). This produces a set of images with no background (0.00), a background similar to what would be acquired in an experiment (0.25-0.75), and an excessive level of background (1.00). We then ran SynBot on these images using manual, ilastik, or SynQuant thresholding. Since the position of these simulated synapses was known, we computed the recall (True Positive / (True Positive + False Negative)) (**Figure 4B**) and precision (True Positive / (True Positive + False Positive)) (**Figure 4C**) for each method on our simulated images. These metrics are well suited to analyses where true positive, false positive, and false negative counts are obtained but true negative counts are not well defined (every point on the image without a synapse could be considered a true negative). We found that the recall and precision values were similar for each method. However, the methods all had decreased accuracy when the images contained excessive background. We found no significant differences between the recall or precision obtained with these thresholding methods when using a 2-way ANOVA at each noise level. Together these analyses indicate that SynBot accurately counts synapses in realistic images containing low and high background noise.

**Figure 4:**
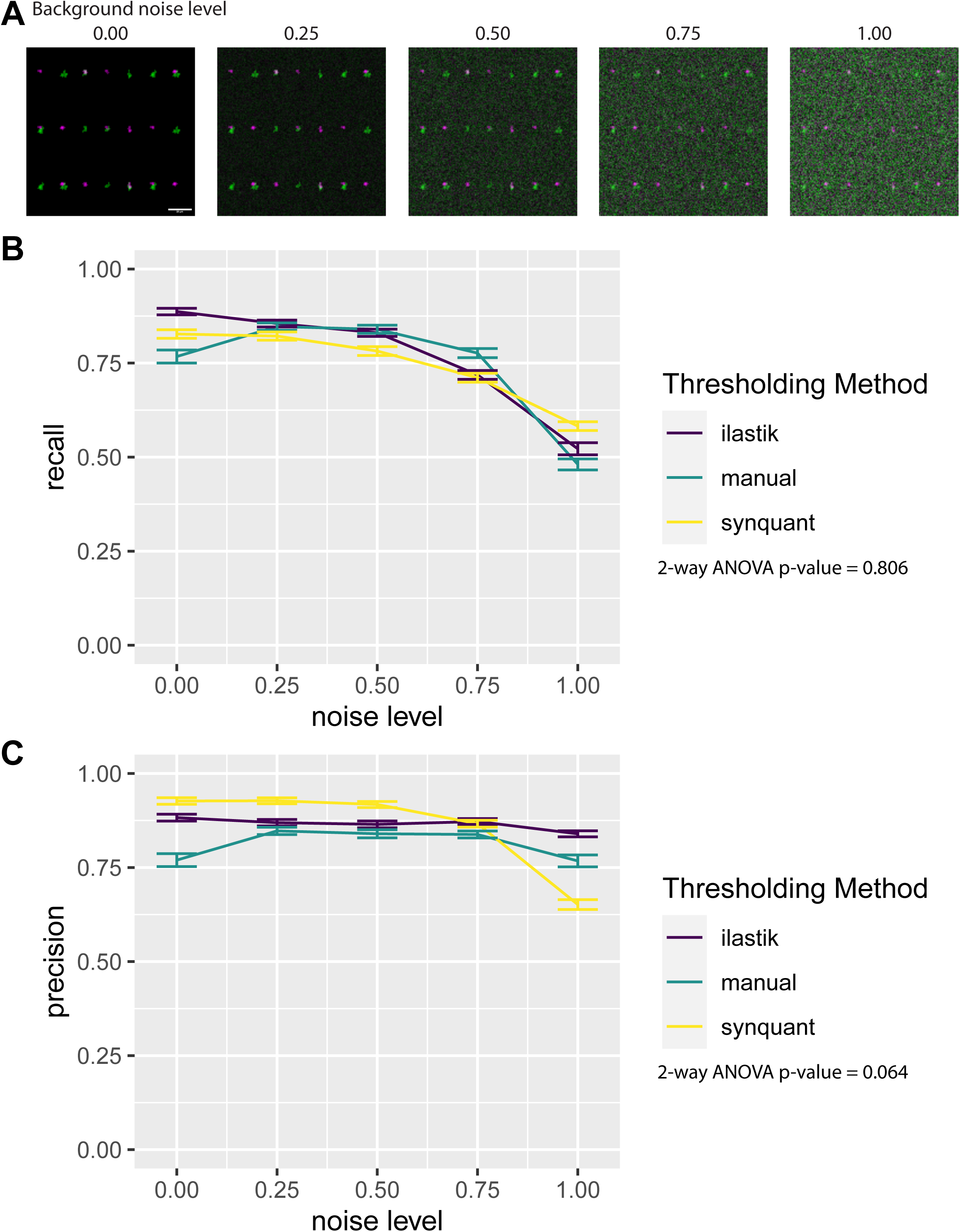
SynBot quantifies simulated synapse data with high recall and precision. a. Simulated synapse images with differing levels of gaussian noise background. Synapses were copied from VGlut1-PSD95 a2d1 WT and KO images from figure 6 and then pasted onto Gaussian noise background with a mean and standard deviation proportional to the same values in the original image multiplied by the given noise level (0.00, 0.25, 0.50, 0.75, 1.00). Error bar represents 20um.
b. Recall (True Positive / (True Positive/ False Negative)) obtained using SynBot with manual, ilastik, and SynQuant thresholding methods. 20 images were analyzed at each noise level. Error bars represent 1 standard error of the mean.
c. Precision (True Positive / (True Positive/ False Negative)) obtained using SynBot with manual, ilastik, and SynQuant thresholding methods. 20 images were analyzed at each noise level. Error bars represent 1 standard error of the mean.

### Quantification of astrocyte-induced synaptogenesis by SynBot *in vitro*

To test SynBot’s utility and accuracy, we next used a glia-free cortical neuronal culture system^61,62^. These neurons form few synapses when cultured alone; however, astrocytes secrete proteins that strongly promote excitatory and inhibitory synapse formation in these cultures^61,62^. The synaptogenic effects of astrocyte-conditioned media (ACM) treatment are robust and promote a highly reproducible increase in the colocalization of pre and postsynaptic puncta. This histological effect has been validated by numerous studies using electron microscopy and electrophysiology^61,63–66^. Therefore, to test the utility of SynBot and compare it to Puncta Analyzer, here we used ACM treatment of purified cortical neuron cultures to induce synapse formation.

For these studies, neurons and astrocytes were individually prepared from postnatal day 1 (P1) Sprague-Dawley rat pups. First, we dissected the cerebral cortex and performed enzymatic digestion with papain and DNAse inhibitor for 45 minutes at 32C. To stop the enzymatic digestion, we treated the tissue with ovomucoid, followed up by mechanical dissociation to obtain a single cell suspension (**Figure 5A**). This single cell suspension was then used to isolate neurons or astrocytes.

**Figure 5:**
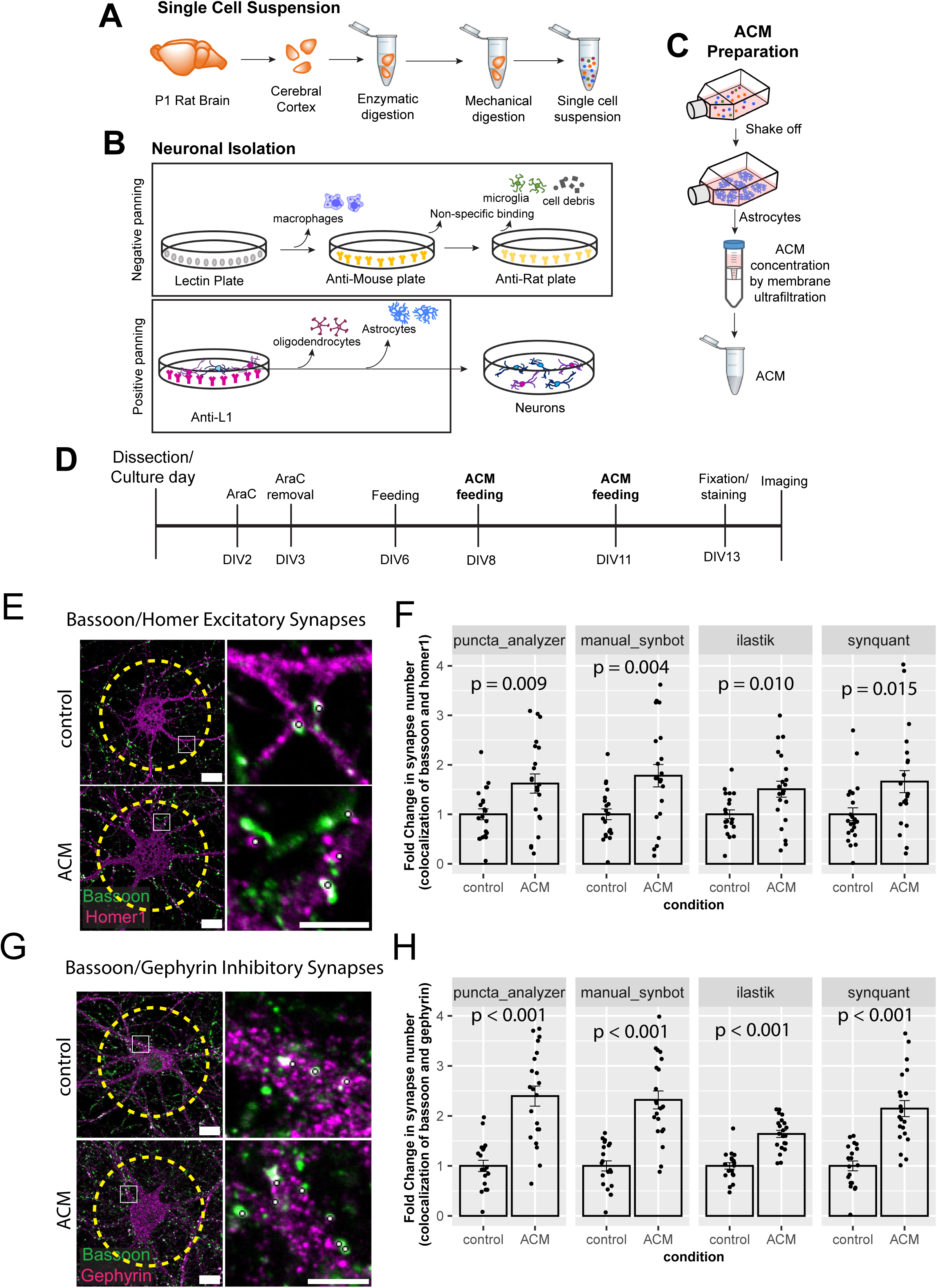
SynBot quantifies astrocyte-induced synapse formation *in vitro*. a. Scheme of single cell suspension generation from P1 rat pup cortices.
b. Scheme of neuronal isolation by immunopanning first through a series of negative panning steps and then through positive panning with Anti-L1CAM antibody.
c. Scheme of astrocyte purification and astrocyte-conditioned media (ACM) preparation.
d. Timeline of neuronal feeding schedule and ACM treatment.
e. Representative images (in RGB) of excitatory synaptic puncta stained with presynaptic marker Bassoon and postsynaptic marker Homer1. Scale: 15 μm for full image and 5 μm for zoom in. Yellow dashed line represents the region of interest analyzed. White dots represent counted synapses.
f. Example of quantification of excitatory synaptic colocalization between control and ACM using the Puncta Analyzer, SynBot manual thresholding, SynBot ilastik, or SynBot SynQuant thresholding. Student’s T-test, 20 neurons per treatment. Error bars represent 1 standard error of the mean.
g. Representative images (in RGB) of inhibitory synaptic puncta stained with presynaptic marker Bassoon and postsynaptic marker Gephyrin. Scale: 15 μm for full image and 5 μm for zoom-in. Yellow dashed line represents the region of interest analyzed. White dots represent counted synapses.
h. Example of quantification of inhibitory synaptic colocalization between control and ACM using the Puncta Analyzer, SynBot manual thresholding, SynBot ilastik thresholding, or SynBot SynQuant thresholding. Student’s T-test, 20 neurons per treatment. Error bars represent 1 standard error of the mean.

As illustrated in **Figure 5B**, neurons were purified from the single cell suspensions first through a series of negative panning steps to remove unwanted cell types in the following manner: first using Baneiraea Simplicifolia Lectin 1, we removed macrophages, second, using an anti-mouse secondary antibody-coated plate and an anti-rat secondary antibody-coated plate we removed potential non-specific Fc-binding cells such as microglia and cell debris. After the negative selection, we then incubated the cell suspension on a positive panning petri dish coated with anti-L1CAM antibody. Neonatal cortical neurons highly express L1CAM, which is absent from other cortical cell types, such as astrocytes and oligodendrocytes. Both excitatory and inhibitory neurons of the cortex are retained on the plate after incubation, and gentle washes are used to remove unbound cells. Finally, the neurons were collected from the positive-panning plate by gentle pipetting and pelleted for resuspension in neuronal growth media and plated on glass coverslips which are coated with poly-D-lysine (PDL) and laminin at a density of 70 thousand cells per coverslip.

To isolate astrocytes, we plated the cortical single-cell suspension onto a PDL-treated tissue culture flask. After 3 days *in vitro* (DIV3), we purified the astrocytes, which are tightly attached to the flask, by vigorously shaking so less-adherent cells are removed. To collect the ACM, we incubated the astrocytes with minimal media for 4 days to promote protein secretion. We collected the ACM and concentrated it using centrifugal concentrator tubes (**Figure 5C**). After protein quantification, we applied 50ug/ml or 100ug/ml ACM to the glia-free neuronal cultures at DIV8 and DIV 11 to induce excitatory and inhibitory synapse formation, respectively (**Figure 5D**).

To label excitatory and inhibitory synapses in culture, we fixed the DIV 13 neuronal cultures and immuno-stained the synapses with pre- and postsynaptic markers. To identify excitatory synapses, we used the presynaptic active zone marker Bassoon together with the excitatory postsynaptic marker Homer1 (**Figure 5E**). To visualize inhibitory synapses, we used Bassoon in combination with the inhibitory postsynapse marker Gephyrin (**Figure 5G**). For image acquisition, we took single focal plane images using an inverted Olympus FV3000 confocal laser scanning microscope with a 60X oil immersion objective. The images were taken in a 1024X1024 resolution with a 2X digital zoom. To identify individual neurons, we used the DAPI channel, and we imaged only the neuronal cell bodies at least two cell diameters distant from other neurons. We used this strategy to avoid overlapping cells and to use an unbiased method to select the neurons to image. We imaged at least 20 cells for each condition, and the laser power for each channel was adjusted such that the signal for each channel was bright without many saturated pixels.

To determine the efficiency of SynBot to detect pre- and postsynaptic colocalization, we analyzed images from cortical neurons treated with ACM or cultured in growth media only (named here as control) using manual, ilastik, and SynQuant thresholding methods of SynBot. We compared these results to analyses done by its predecessor Puncta Analyzer (**Figure 5E-H and S2**). To aid these comparisons, we normalized the number of excitatory synapses between the ACM-treated neurons to the control values. As expected, ACM induced a significant increase in synapse numbers (∼1.5-fold) with each of the analysis methods used (student’s t test p = 0.005, p = 0.004, p = 0.010, and p = 0.0150 for Puncta Analyzer, SynBot Manual, SynBot ilastik, and SynBot SynQuant respectively) (**Figure 5F**). Similarly, we found a 2-fold increase in the number of inhibitory synapses when neurons were treated with ACM compared to the control (student’s t-test p < 0.001 for each analysis method) (**Figure 5H**). These results indicate that SynBot is equally efficient in detecting both types of synaptic contacts as the Puncta Analyzer. Moreover, these results show that the ilastik and SynQuant automated thresholding methods can be used to determine changes in synapse numbers.

To determine whether manual or automated thresholding affected the numbers of puncta identified, we also compared the raw numbers of green (presynaptic marker), red (postsynaptic marker), and colocalizations identified between the Puncta Analyzer, SynBot Manual, SynBot ilastik, and SynBot SynQuant. When compared to the other methods, SynBot ilastik consistently identified more green, red, and colocalized puncta from the images, regardless of the antibodies used or the treatment conditions (**Figure S1**). This result is consistent with ilastik’s tendency to detect more individual objects (here puncta) based on 37 different image features. In contrast, manual thresholding only uses intensity cutoffs and minimal puncta size as features for detection. Two-way ANOVA tests found significant differences in the raw number of colocalized synapses detected by each of the analysis methods for the *in vitro* excitatory and inhibitory datasets shown in **Figure 5**. When the number of colocalized puncta detected was normalized to the appropriate control conditions; however, there were no significant differences between the 4 thresholding methods. Together, these results show that SynBot, with its user-friendly and timesaving features, efficiently detects excitatory and inhibitory synaptic contacts *in vitro*.

### Quantification of *in vivo* excitatory synapse numbers using SynBot

After validating the use of SynBot *in vitro*, we next tested its utility in quantifying synapse numbers from mouse brain tissue sections. Immunohistochemistry of brain tissue sections is a common method for synapse number quantification^40,41,45,46,57,62^. However, these analyses have the considerable added complexity of a 3-dimensional tissue section that should be imaged. Moreover, thick tissue sections from the brain inherently have more background signals, which can impair the accuracy of the analyses. Finally, antibody penetration into the tissue sections is an important consideration while optimizing synapse staining and imaging procedures. Before using SynBot or any other method, the experimenters should optimize their staining and imaging procedures. The optimized synapse staining and imaging procedure we have used can be found in the methods and protocols.io (https://www.protocols.io/view/synbot-protocols-3byl4qewjvo5/v2).

To test SynBot’s efficacy and accuracy in quantifying synapse numbers *in vivo*, we next analyzed images that were previously acquired and reported in a study by Risher and colleagues in 2018^57^. This paper showed that the neuronal Thrombospondin/Gabapentin receptor α2δ-1 is crucial for proper excitatory synapse formation in the developing mouse cortex^57,64,67^. Risher et al., 2018 showed that in α2δ-1 KO animals, there is a significant decrease in VGluT1/PSD95 synapse numbers compared to WT controls^57^. Risher et al. validated these histological findings of impaired synaptogenesis using electrophysiology and electron microscopy that also showed a strong decrease in excitatory synapse function and number, respectively.

To perform these experiments, mice were transcardially perfused with 4% paraformaldehyde, and brains were collected, frozen, and cryosectioned (**Figure 6A**). Intracortical excitatory synapses were marked with the pre- and post-synaptic markers specific for these connections, namely vesicular glutamate transporter 1 (VGluT1) and postsynaptic density 95 (PSD95) (**Figure 6B**). Images were acquired using a Leica SP5 confocal microscope. A z-stack of 15 optical planes was acquired using a 0.34 um z-step to image ∼ 5 μm of the sample. Images were acquired such that confocal settings were optimized on control conditions to minimize oversaturated pixels. For detailed procedures, see the methods section and protocols.io (https://www.protocols.io/view/synbot-protocols-3byl4qewjvo5/v2).

**Figure 6:**
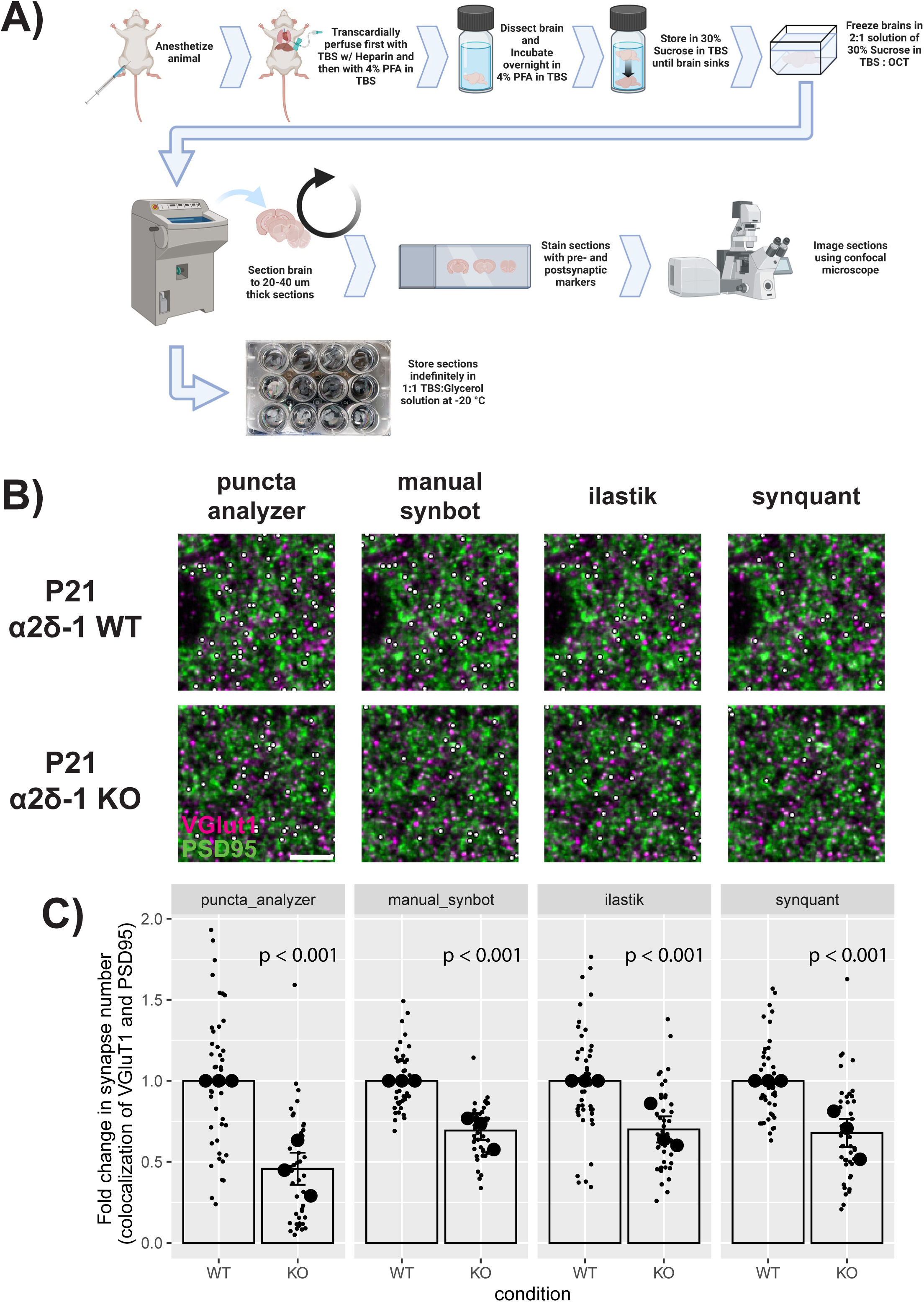
SynBot quantifies reduced α2δ-1 KO synapse numbers *in vivo*. a. Workflow for harvesting mouse brain tissue and preparing it for synapse imaging.
b. Representative images of synapses from layer 2/3 of primary visual cortex in α2δ-1 wild type vs. α2δ-1 knockout mice at postnatal day 21. Magenta = VGluT1; Green = PSD95. Synapses identified by each thresholding method are marked by small white dots. Scale bar represents 5 μm.
c. Quantification of VGluT1-PSD95 colocalization using the Puncta Analyzer, SynBot manual thresholding, SynBot ilastik thresholding, or SynBot SynQuant thresholding. Colocalized puncta counts were normalized to the WT average for each experimental pair. A linear mixed-effects model was used to account for multiple images per animal. Mouse averages are shown as large black dots (N = 3 mice per condition) with individual images shown as small gray dots. Error bars represent 1 standard error of the mean.

We used this well-validated data set to test SynBot’s efficacy for quantifying synapse numbers in mouse brain tissue images. We reanalyzed the P21 α2δ-1 WT and KO images previously published in Risher et al., 2018 using the SynBot manual, ilastik, and SynQuant thresholding methods and compared these to the results Risher et al. obtained using Puncta Analyzer (**Figure 6C**). All 4 methods found a significant ∼50% decrease in the number of synapses in α2δ-1 KO animals when compared to WT littermates (linear mixed effects model, p < 0.001 for each method). In our hands, a pre-trained ilastik model implemented in SynBot or SynQuant automated thresholding was able to perform as well as a highly trained manual user (**Figures 6C and S1)**. As we saw with the *in vitro* datasets, there were significant differences between the raw numbers of colocalized puncta detected by the 4 thresholding methods but not after normalization to the WT control data (two-way ANOVA with p-value cutoff of 0.05). Quantification using SynBot manual, ilastik, or SynQuant thresholding recapitulated the previously validated finding that α2δ-1 KO mice have fewer intracortical synapses than WT littermates, which was also previously confirmed by EM and electrophysiology^57^. This demonstrates that SynBot can be successfully applied to brain tissue images and that the thresholding methods packaged in SynBot can be used to compare different strategies quickly.

Altogether SynBot is a fast, reliable, user-friendly, and versatile software to study synapse density *in vitro* and *in vivo*, overcoming previous constraints of Puncta Analyzer.

## Discussion

Quantification of synapse numbers using immunohistochemistry is a favored technique for its ability to label synapses quickly and easily. However, this rapid experimental protocol is often paired with slow analysis methods that are difficult to learn and result in variability between users. SynBot overcomes many of the shortcomings of its predecessor by being easy to learn with built-in and online instructions and easy to use with a user interface that simplifies and streamlines user input. SynBot also allows unsupervised analyses through its automated methods. Importantly, SynBot collects all the data extracted from synaptic images by recording the position and area of each individual punctum from each image channel. These data can be used for other forms of analyses, such as the count and sizes of individual synaptic puncta or the spatial relationship in their distribution across the tissue imaged.

One of the most challenging aspects of analyzing these data is the subjective nature of the manual thresholding used in the Puncta Analyzer. The 9 thresholding modes available in SynBot address this by providing the user with simple approaches like a fixed threshold value, streamlined manual thresholding as well as the complex machine learning-based algorithm of ilastik and probability-based algorithm of SynQuant. We have found ilastik to be particularly suited to this workflow as a short training session in ilastik is sufficient to analyze images accurately and reproducibly without the need for further user input. The details about how to train ilastik for synaptic puncta detection can be found in the methods section and protocols.io. When used in the manual mode, SynBot also automatically saves the threshold values applied to each image with the output data, allowing for rapid replication of the analysis by other users.

The advantage of implementing SynBot in the ImageJ macro language is the ease with which it can be changed by users to apply to a wider variety of experimental questions. All the source code for the macro is available on our lab GitHub site (https://github.com/Eroglu-Lab/Syn_Bot) and can be edited within FIJI itself without the need for software development programs or in-depth computer science experience. There were only a few parts of SynBot that could not be implemented in ImageJ macro language (circular-approximation colocalization, some dialog menus, ilastik and SynQuant integration) and required Java programming. These sections are stored in the separate GitHub repositories ilastik4ij_Syn_Bot (https://github.com/Eroglu-Lab/ilastik4ij_Syn_Bot) and (https://github.com/freemanwyz/SynQuantSimple), which are freely available for download and can be edited by users with experience in Java programming.

Another benefit of SynBot over previous analysis programs is its speed. When analyzing the 100 simulated images shown in Figure 4, for example, SynBot analysis took 24 minutes for manual thresholding, 10 minutes for training and 50 minutes for ilastik thresholding, and 4 minutes with SynQuant thresholding. Unlike manual thresholding, the time spent while ilastik or SynQuant run requires no user input. The speed of SynQuant makes it well-suited for running multiple times using different parameters, making it easy to troubleshoot and optimize accuracy for a given data set.

With all of these features, researchers will be able to rapidly screen experimental conditions that alter synapse numbers and can move on to in-depth structural and mechanistic investigations with other methods such as electrophysiology, super-resolution or electron microscopy. Therefore, SynBot will aid in answering many outstanding questions related to synapse development and maintenance.

### Limitations of the Study

There are a few key conditions necessary for SynBot to be successfully applied. Firstly, the signals being analyzed need to be punctate rather than diffuse since SynBot performs object-based colocalizations and must be able to identify discrete objects. Similarly, objects within the images must be non-overlapping to be independently counted. This object-based colocalization is most appropriate when the position of synaptic compartments is changing to alter the number of structural synapses (seen as colocalizations). If the abundance of a synaptic marker or its association with synapses is changing, intensity measurements or pixel-wise colocalization on unthresholded raw images may be more appropriate (see https://imagej.net/imaging/colocalization-analysis for further discussion of different colocalization methods).

SynBot uses synaptic puncta position over raw intensity. Thus, we can use thresholding methods such as ilastik and SynQuant that apply different values to each image to separate the foreground from the background. However, if needed SynBot can also apply the “Fixed Value” thresholding, which allows users to apply the same exact threshold value to all images. An important limitation of this fixed thresholding method is that it would only be useful for datasets with images containing minimal background noise, such as *in vitro* synapse staining.

While theoretically possible, in practice staining intensities and background may vary even within a single experiment enough to impair the use of fixed values. We strongly recommend using methods like ilastik or SynQuant instead because they have the flexibility to adjust to the varying image brightness. In particular SynQuant thresholds images using a statistical method that provides consistency between different images. Thus, these automated methods make these analyses much more accessible for untrained new users than manual thresholding.

SynBot relies on RGB image conversion to keep track of the different image color channels. This limits SynBot analysis to a maximum of 3 colors and converts each image channel to an 8-bit format. This reduction in bit-depth from the 16-bit images acquired by many modern microscopes reduces the possible thresholding values from 65,536 in a 16-bit image to 256 in an 8-bit image. For the synapse analysis experiments shown here, 256 possible values is adequate for thresholding, but certain analyses may benefit from dividing the image intensity range into a greater number of bins. It should be noted that SynBot has no limitations for image resolution (the number of pixels representing a given area of space) and that this bit-depth conversion has no impact on resolution.

SynBot only works on single optical sections or Z-stack projections and is unable to fully incorporate 3-dimensional synapse structures, as is done in other software such as Imaris. Despite these caveats, SynBot is widely applicable to any object-based colocalization analysis for synapses or any other set of imaging markers. SynBot can tailor its analysis to many experimental questions. However, for neurite tracing synapse analysis or 3D constructions alternative methods should be used^52,53^. Similarly, if the user needs to analyze multiple regions from an image, they should perform repetitive analysis to follow that experimental design.

It is also important to note that the accuracy of synapse number analyses is heavily dependent on the quality of staining and imaging. Moreover, proper markers should be used to label synapses. Some common mistakes are to use an axonal or dendritic marker, such as GAD65 or GAD67 (for inhibitory axons) or MAP2 (for neuronal dendrites), as one of the synaptic compartments. These kinds of markers are often extrasynaptically localized and widely distributed, thus yield co-localization with other markers even if they were not at an actual synapse.

Similarly, SynBot requires careful troubleshooting, even when using fully-automated thresholding methods. Users should always visually inspect the counted synapses using the “colocs” output images and verify the thresholding performance by viewing the “red_thresholded” and “green_thresholded” output images. It is also advisable to include control conditions whenever possible, such as perturbations known to change synapse numbers. As with any image analysis method, improper use can lead to misleading results.

## Supporting information

Key Resources

## Acknowledgments

We thank Christabel Tan, M. Pia Rodriguez-Salazar, Dr. Oluwadamilola Lawal, Nicholas Brose, Gabrielle Sejourne, Kavya Raghunathan and other Eroglu lab members for critical feedback about the manuscript. This work was supported NIH R01 funding AG059409, NS102237, and BRAIN Initiative funding U19NS123719 and a Chan Zuckerberg Initiative, Neurodegeneration Challenge Network Collaborative grant (2018-191999 and DAF2021-237435) and by the joint efforts of MJFF and the Aligning Science Across Parkinson’s (ASAP) initiative. MJFF administers the grant [ASAP-020607 to CE] on behalf of ASAP and itself Michael J Fox Foundation. J.T.S. was supported by NIH funding F31NS134252. D.I. was supported by The Pew Latin American Fellows Program in the Biomedical Sciences, Foerster-Bernstein Post-Doctoral Fellowship Program for Women in STEM, and the Holland-Trice Scholars Award for high risk/high impact discovery in research in Basic Brain Science and Brain Diseases. JJR was supported by NIH funding NS102237-S1 and F31NS125985. YW was supported by NIH funding R01MH110504. WCR was supported by NIH funding NIH R15MH126345. Dr. Cagla Eroglu is an HHMI Investigator. The cartoon of a synapse in Figure 1 and a diagram of mouse brain tissue staining in Figure 5 were created using BioRender.com.

## IP rights notice

This article is subject to HHMI’s Open Access to Publications policy. HHMI lab heads have previously granted a nonexclusive CC BY 4.0 license to the public and a sublicensable license to HHMI in their research articles. Pursuant to those licenses, the author-accepted manuscript of this article can be made freely available under a CC BY 4.0 license immediately upon publication.

## Author Contributions

Conceptualization, J.T.S. and C.E.; Software, J.T.S., J.R., Y.W.; Investigation, J.R., W.C.R., and D.I.; Writing – Original Draft, J.T.S., J.R., D.I., and C.E.; Writing – Review & Editing, J.T.S., J.R., D.I., and C.E.

## Methods

### RESOURCE AVAILABILITY

#### Lead contact

Further information and requests for resources and reagents should be directed to and will be fulfilled by the lead contact, Cagla Eroglu.

#### Materials availability

This study did not generate new unique reagents.

#### Data and code availability

- Microscopy data and associated synapse counts reported in this paper are available on Zenodo at the following DOI: 10.5281/zenodo.12191805
- All original code has been deposited at our lab Github site (https://github.com/Eroglu-Lab/Syn_Bot) and is publicly available on Zenodo at the following DOI: 10.5281/zenodo.12192447
- Any additional information required to reanalyze the data reported in this paper is available from the lead contact upon request.

## EXPERIMENTAL MODEL AND SUBJECT DETAILS

### Animals

Mice and rats were used for experiments as specified in the text and figure legends. All mice and rats were used in accordance with the Institutional Animal Care and Use Committee (IACUC) and the Duke Division of Laboratory Animal Resources (DLAR) oversight (IACUC Protocol Numbers A147-17-06 and A117-20-05). All mice and rats were housed under typical day/night conditions of 12-hour cycles. For all experiments, age, and sex-matched mice were randomly assigned to experimental groups based on genotypes. The primary rat neurons and astrocytes were isolated from wildtype Crl: CD(SD) Sprague-Dawley rats from Charles River Laboratories (RRID: RGD_734476).

## METHOD DETAILS

### In vitro Synapse Assay with Astrocyte Conditioned Media

#### Cortical tissue digestion

Primary rat astrocytes and neurons were isolated from postnatal day 1 (P1) rat pups. Pups were rapidly decapitated, and brains dissected out and placed into a petri dish of PBS. Brains were then micro-dissected to isolate the cerebral cortex, and the cortices were chopped into pieces approximately 1 cubic millimeter in volume. A transfer pipet was then cut where it began to taper to create a larger opening. This cut pipet was then used to suck up the cortex chunks, and tissue was transferred to a tube of 7.5 units/mL papain digestion solution (Worthington, Cat# LK003178) by gently tapping on the pipette without depressing it. Cortices were then digested for 45 minutes at 33°C with a brief swirling at the 15- and 30-minute points. The digestion solution is then aspirated off and the tissue is resuspended in a 2 mg/ml trypsin inhibitor ovomucoid solution (Lo Ovo) (Worthington, Cat# LS003083). Cells were then spun in a centrifuge at 300g for 11 minutes at room temperature. The supernatant was then removed, and the cells were resuspended in a 4 mg/ml Ovomucoid solution (Hi Ovo). Cells were then spun again at 300g for 11 minutes and the supernatant removed. The resulting cells were then either used for isolating neurons or astrocytes according to the following sections.

#### Neuronal Isolation by Immunopanning

The pelleted cells from the cortical tissue digestion were resuspended in Panning Buffer (0.02% BSA with 0.5 ug/ml insulin in DPBS with calcium, magnesium, glucose, and pyruvate (Thermo Fisher, Cat# 14287080)). The cells were then filtered through a 20μm nylon mesh (Elko Filtering, Cat# 03-20/14) to remove clumps. These filtered cells were then added to a petri dish which had been treated with Griffonia Simplicifolia Lectin I (Vector Laboratories, Cat# L-1100) for 1 hour at room temperature prior to adding the cells. The cells were incubated on this lectin plate for 10 minutes at room temperature. The plate was then forcefully shaken to resuspend any loosely adhered cells. The solution in the lectin plate was then transferred to a second lectin plate for a 15-minute incubation at room temperature. The second lectin plate was then forcefully shaken, and the solution transferred to a plate treated with an L1CAM antibody (α-L1) (Developmental Studies Hybridoma Bank, Cat# ASCS4) which binds to neurons. The cells were incubated on the α-L1 plate for 45 minutes at room temperature. The media was then gently removed from the α-L1 plate and the plate was washed 5 times with panning buffer. Cells bound to the α-L1 plate were then resuspended in the panning buffer and collected in a conical tube. The cells were then spun for 11 minutes at 300g. The supernatant was removed, and the cells were resuspended in neuronal growth media (NGM: Neurobasal (Gibco, Cat# 21103049), B27 supplement (Gibco, Cat# 17504044), 2 mM L-Glutamine (Gibco, Cat# 25030-081), 100 U/mL Pen/Strep (Gibco, Cat# 15140), 1 mM sodium pyruvate (Gibco, Cat# 11360-070), 4.2 μg/mL Forskolin (Sigma, Cat# F6886), 50 ng/mL BDNF (PeproTech, Cat# 450-02), and 10 ng/mL CNTF (PeproTech, Cat# 450-13)). The cells were then counted, and plated at a low density of 60,000 cells each onto poly-D-lysine-treated (Sigma, Cat# P6407) and mouse laminin (Cultrex, Cat# 3400-010-01) treated glass coverslips in a 24-well plate with neuronal growth media. Cells were then kept in a tissue culture incubator at 37°C with 10% CO_2_.

#### Astrocyte Isolation and ACM Production

The pelleted cells from the cortical tissue digestion were resuspended in astrocyte growth media (AGM: DMEM (GIBCO, Cat# 11960), 10% FBS (Thermo Fisher, Cat# 10-437-028), 10 mM, hydrocortisone (Sigma, Cat# H-0888), 100 U/mL Pen/Strep, 2 mM L-Glutamine, 5 mg/mL Insulin (Sigma, Cat# 11882), 1 mM Sodium Pyruvate, 5 mg/mL N-Acetyl-L-cysteine (Sigma, Cat# A8199)). The cells were then filtered through a 20μm nylon mesh to remove cell clumps. The filtered cells were then spun for 9 minutes at 300g. The supernatant was then removed, and the cells were resuspended in AGM and counted. 15 million cells were then plated in a T-75 flask with a non-aerating top. Cells were then kept in a tissue culture incubator at 37°C with 10% CO_2_ with the cap slightly unscrewed to allow airflow.

On DIV3, astrocyte flasks were washed 3 times with DPBS without calcium or magnesium (Gibco, Cat# 14190144). After the third wash, 20 mL of DPBS without calcium or magnesium was added and the flask was vigorously shaken to remove loosely adherent cells. This shake-off process significantly enriches for astrocytes. Media was then replaced with AGM and flasks were returned to their incubator. On DIV 5, astrocytes were treated with 2.43 mg/ml Cytosine arabinoside (AraC) (Sigma, Cat# C1768) through a full media change. On DIV 7 astrocytes were passaged using 0.05% Trypsin-EDTA. 3 million cells each were then plated into 10 cm culture dishes. On DIV 9, astrocytes were switched to a minimal media (Neurobasal medium minus phenol red (Gibco, Cat# 12348017), 100 U/mL pen/strep, 2mM L-glutamine, and 1mM sodium pyruvate) for astrocyte conditioning. Astrocytes were then cultured for 5 days without any media change, and on DIV 14, the ACM was collected. The ACM was then centrifuged for 5 minutes at 1,100g to pellet cellular debris. The supernatant was concentrated by centrifugation for 1 hour at 3220 g at 4°C in a 5kDa Cutoff Vivaspin tube (Sartorius, Cat # VS2012). ACM was aliquoted into low protein binding tubes (∼100ul per aliquot), flash-frozen in liquid nitrogen, and stored at −80°C until use. An aliquot of each batch of ACM was thawed and used to measure the protein content using the Pierce BCA protein assay kit (Thermo Fisher, Cat# 23225).

#### Neuron Feeding and ACM Treatment

On DIV2, neurons were treated with 2.43 mg/ml AraC through a half-media change. A full media change was then performed 24 hours later (DIV3) to remove the AraC. Neurons were then given another half-media change on DIV6. Neurons were then fed with ACM on DIV8 and DIV 11 by adding either 50 mg/ml ACM for excitatory synapses or 100 mg/ml ACM for inhibitory synapses in a half media change.

#### Neuron Staining and Imaging

On DIV 13, neurons were fixed with 4% paraformaldehyde (Electron Microscopy Sciences, Cat# 15710) in PBS. Cells were then washed 3 times with PBS and then blocked for 30 minutes at room temperature with antibody blocking buffer (150 mM NaCl, 50 mM Tris-Base, 1% BSA, 100 mM L-lysine 50% normal goat serum (Thermo Fisher, Cat# 01-6201), 0.2% Triton (Roche, Cat# 11332481001). The blocking buffer was then removed, and cells were incubated with primary antibodies against the pre- and post-synaptic markers of interest overnight at 4°C in antibody incubation buffer (150 mM NaCl, 50 mM Tris-Base (VWR, Cat# 101174-856), 1% BSA, 100 mM L-lysine, 10% normal goat serum). The following day cells were washed 3 times with PBS and incubated in fluorescent secondary antibodies in antibody incubation buffer for 2 hours at room temperature protected from light. Cells were then washed with PBS 3 times and incubated with 1:10,000 DAPI (Invitrogen, Cat#D1306) for 5 minutes. Cells were then washed 3 times with PBS and mounted onto coverslips with homemade mounting media (20mM Tris pH 8.0, 90% Glycerol, 0.5% N-propyl gallate) and sealed with nail polish.

Single focal plane images were taken using an inverted Olympus FV3000 confocal laser scanning microscope with a 60X oil immersion objective. The images were taken in a 1024X1024 resolution with a 2X digital zoom. For the picture acquisition, the DAPI channel was used to select neuronal cell bodies distant from other neurons to avoid overlap of cell soma and to do a blind selection of the neurons. DAPI positive nuclei with abnormal morphology were excluded for the analysis. 20 cells for each condition were selected for the analysis. Images were also saved as .bmp extension with only Bassoon (green) and Gephyrin (red channel) for the synaptic analysis.

### Mouse Brain Tissue Synapse Assay

#### Mouse perfusion and cryosectioning

P21 α2δ-1 WT and KO mice were euthanized by intraperitoneal injection of 0.6mL of 12.5 mg/mL 2,2,2-tribromoethanol (Avertin) (Sigma, Cat# T48402-25G) followed by exsanguination through transcardially perfusion of tris-buffered saline (TBS: 137mM NaCl, 2.68 mM KCl, 24.8 mM Tris-base (Thermo Fisher, Cat# J75825.A7)). Following TBS perfusion, animals were perfused with warm, 4% paraformaldehyde (PFA) in TBS solution. Brains were then removed and post-fixed in 4% PFA overnight at 4°C. After post-fixation, brains were washed 3 times with TBS and then incubated in 30% sucrose in TBS for 48-72 hours. Brains are fully cryopreserved once they sink to the bottom of their container in the 30% sucrose. Brains were then placed in cryomolds and frozen in 50% Optimal Cutting Temperature Compound (OCT) (Tissue Tek, Cat# 4583) in TBS on dry ice. Brains were then kept at −80°C until sectioning. Brains were cryosectioned to a thickness of 15-40 μm and stored in 50% glycerol (Acros Organics, Cat# 15892-0010) in TBS at −20°C.

#### Tissue Section Synapse Staining and Imaging

Three tissue sections per animal were washed 3 times for ten minutes with 0.2% TritonX-100 in TBS (TBST) on a shaker at room temperature. Sections were then blocked with 5% normal goat serum in TBST for 1 hour at room temperature on a shaker. Sections were then stained with primary antibodies for the pre and postsynaptic markers of interest overnight at 4°C on a shaker. Sections were then washed 3 times in TBST for 10 minutes and incubated with fluorescent secondary antibodies for 2 hours at room temperature on a shaker protected from light. After secondary incubation, sections were washed 3 times with TBST for 10 minutes. After washing, sections were mounted onto microscope slides and sealed with nail polish.

Stained tissue is ready to image after nail polish is dried and should be imaged within 48 hours of secondary staining for most accurate results. Images were acquired using a Leica SP5 confocal microscope with a 63X oil immersion objective. For 1024 × 1024 images, a 1.64X digital zoom was used to achieve an XY resolution of 0.126 um X 0.126 μm pixel size and a z-stack of 15 optical planes was acquired using a 0.34 μm z-step to image ∼ 5 μm of the sample. In general, images were acquired such that confocal settings were optimized on control conditions with the goal of minimizing oversaturated pixels but maintaining real synaptic puncta.

### Image Analysis

#### Synapse Quantification with Puncta Analyzer

*In vitro* synapse assay images were analyzed using Puncta Analyer^38^ by first converting the images to the RGB format and then running the Puncta Analyzer plugin on each image. A circular ROI was applied to each image with a radius of 301 pixels around the cell body of the imaged neuron. A threshold for each image was chosen to minimize the background while preserving the signal.

*In vivo*, synapse assay images were Z-projected such that every 3 optical planes were projected to produce 5 max-projections corresponding to 1 μm, per 15-image stack covering the 5 μm. Images were then analyzed with the Puncta Analyzer plugin using default settings. A threshold for each image was chosen by users to minimize the background while preserving the signal.

#### Synapse Quantification with SynBot

Images were analyzed with the SynBot macro using the default settings for the manual thresholding method. A threshold for each image was chosen to minimize the background while preserving the signal. Threshold performance was checked visually using the “_colocs” image SynBot creates that displays which puncta were counted as synapses in each image.

For the ilastik thresholding method, images were first preprocessed into individual color channels using the extract_channels macro (available at https://github.com/Eroglu-Lab/Syn_Bot). These single-channel images were then used to train ilastik projects for each image channel. Each ilastik project was trained using the pixel classification mode in ilastik. A random subset of images from the target color channel was chosen for the training set. For both *in vitro* and *in vivo* synapse analysis, we used all the image features available. We then annotated approximately 20 synapses as label 1 and marked several regions of the background as label 2 on the first image. We then used the live update feature to view the classifications of each pixel and added additional annotations to the first image and subsequent images as necessary. We then saved the ilastik project files and used these for the SynBot ilastik thresholding method (all ilastik project files used for this paper are available at https://github.com/Eroglu-Lab/Syn_Bot).

For the SynQuant thresholding method, images were analyzed using the SynQuant batch thresholding option and SynBot noise reduction. The following SynQuant parameters were used: Simulated data: Z-score threshold = 10, minimum object size =10, maximum object size = 100, minimum object fill = 0.5, maximum width to height ratio = 4, Z axis multiplier = 1, and estimated noise standard deviation = 12. *In vitro* data: Z-score threshold = 10, minimum object size =10, maximum object size = 100, minimum object fill = 0.5, maximum width to height ratio = 4, Z axis multiplier = 1, and estimated noise standard deviation = 20. *In vivo* data: Z-score threshold = 10, minimum object size =10, maximum object size = 100, minimum object fill = 0.5, maximum width to height ratio = 4, Z axis multiplier = 1, and estimated noise standard deviation = 12.

#### Creation of Simulated Synapse Images

To produce these simulated images, we first used a set of 20 real VGlut1-PSD95 synapse images from Risher et. al., 2019. We split the two image channels and then measured the pixel intensity histogram from these images. We generated a Gaussian noise background for each image by multiplying the mean and standard deviation of the original image’s histogram by a multiplier (0.00, 0.25, 0.50, 0.75, or 1.00). Since the pixel intensity histogram includes the true signal as well as the background, the 1.00 multiplier will produce an image with a higher background than that of the original image. This makes the range of 0 to 1 for our noise multiplier represent images with no background at (0.00), moderate background comparable to experimental data (0.25-0.75), and high background beyond what would be acceptable for analysis (1.00). We next took 10 synaptic puncta from each channel of the original image and pasted these onto the background images we generated. Synaptic puncta were pasted into 1000 possible positions such that 334 positions had a red puncta only, 333 positions had a green puncta only, and 333 positions had a red puncta and a green puncta at the same location. Note that the locations for pasting these synaptic puncta were defined by the top left corner of their bounding box. This resulted in varying levels of overlap similar to real synaptic imaging data and explains the precision and recall of SynBot being below 1.0 since many of these synapses were no longer overlapping after being thresholded. The end product of this code was 100 total images with 20 images having each of the 5 background levels. The ImageJ macro code used for generating these simulated images is available at https://github.com/Eroglu-Lab/Syn_Bot).

## QUANTIFICATION AND STATISTICAL ANALYSES

SynBot was used to quantify synapse numbers in the experiments shown. Subsequent statistical analysis was performed with the R statistical software (R Core Team). Student’s t-tests were used to test for significance between conditions for the *in vitro* experiments, and linear mixed-effects models were used (R package nlme, R Core Team) to test the significance of *in vivo* experiments and account for the nested structure of the data (multiple z-stacks per image and multiple images per animal). Statistical details of each experiment can be found in the figure legends. For the cell culture experiments, each n represents a neuron that was imaged. For mouse tissue sections experiments, each n represents the average synapse density of a mouse that was included in the experiment. For each mouse, at least 15 max-projection images corresponding to 3 independent z-stacks were used. The full R code for performing the statistical analyses and producing the plots in this paper is available at https://github.com/Eroglu-Lab/Syn_Bot.

## References

1. Farhy-Tselnicker, I. & Allen, N. J. Astrocytes, neurons, synapses: a tripartite view on cortical circuit development. Neural Dev 13, (2018).

2. Cadwell, C. R., Bhaduri, A., Mostajo-Radji, M. A., Keefe, M. G. & Nowakowski, T. J. Development and Arealization of the Cerebral Cortex. Neuron 103, 980–1004 (2019).

3. Pan, Y. & Monje, M. Activity Shapes Neural Circuit Form and Function: A Historical Perspective. Journal of Neuroscience 40, 944–954 (2020).

4. Luo, L. Architectures of Neuronal Circuits. Science 373, eabg7285 (2021).

5. Batool, S. et al. Synapse formation: From cellular and molecular mechanisms to neurodevelopmental and neurodegenerative disorders. J Neurophysiol 121, 1381–1397 (2019).

6. Rizalar, F. S., Roosen, D. A. & Haucke, V. A Presynaptic Perspective on Transport and Assembly Mechanisms for Synapse Formation. Neuron 109, 27–41 (2021).

7. Groc, L. & Choquet, D. Linking glutamate receptor movements and synapse function. Science (1979) 368, (2020).

8. Newschaffer, C. J. et al. An Overview of the Main Genetic, Epigenetic and Environmental Factors Involved in Autism Spectrum Disorder Focusing on Synaptic Activity. International Journal of Molecular Sciences 2020, Vol. 21, Page 8290 21, 8290 (2020).

9. Pocklington, A. J., O’Donovan, M. & Owen, M. J. The synapse in schizophrenia. European Journal of Neuroscience 39, 1059–1067 (2014).

10. Lane-Donovan, C. & Herz, J. ApoE, ApoE Receptors, and the Synapse in Alzheimer’s Disease. Trends in Endocrinology & Metabolism 28, 273–284 (2017).

11. Rizo, J. & Xu, J. The Synaptic Vesicle Release Machinery. Annu Rev Biophys 44, 339–367 (2015).

12. Rusakov, D. A., Savtchenko, L. P., Zheng, K. & Henley, J. M. Shaping the synaptic signal: molecular mobility inside and outside the cleft. Trends Neurosci 34, 359–369 (2011).

13. Luján, R., Shigemoto, R. & López-Bendito, G. Glutamate and GABA receptor signalling in the developing brain. Neuroscience 130, 567–580 (2005).

14. Vyklicky, V. et al. Structure, function, and pharmacology of NMDA receptor channels. Physiol Res 63, (2014).

15. Diering, G. H. & Huganir, R. L. The AMPA Receptor Code of Synaptic Plasticity. Neuron 100, 314–329 (2018).

16. Chen, H., Tang, A. H. & Blanpied, T. A. Subsynaptic spatial organization as a regulator of synaptic strength and plasticity. Curr Opin Neurobiol 51, 147–153 (2018).

17. Iasevoli, F., Tomasetti, C. & de Bartolomeis, A. Scaffolding proteins of the post-synaptic density contribute to synaptic plasticity by regulating receptor localization and distribution: relevance for neuropsychiatric diseases. Neurochem Res 38, 1–22 (2013).

18. Akin, O. & Zipursky, S. L. Activity regulates brain development in the fly. Curr Opin Genet Dev 65, 8–13 (2020).

19. Andreae, L. C. & Burrone, J. The role of neuronal activity and transmitter release on synapse formation. Curr Opin Neurobiol 27, 47–52 (2014).

20. Chen, L. F., Zhou, A. S. & West, A. E. Transcribing the connectome: roles for transcription factors and chromatin regulators in activity-dependent synapse development. J Neurophysiol 118, 755–770 (2017).

21. Shen, K. & Scheiffele, P. Genetics and cell biology of building specific synaptic connectivity. Annu Rev Neurosci 33, 473–507 (2010).

22. Sanes, J. R. & Zipursky, S. L. Synaptic Specificity, Recognition Molecules, and Assembly of Neural Circuits. Cell 181, 536–556 (2020).

23. Jain, S. & Zipursky, S. L. Temporal control of neuronal wiring. Semin Cell Dev Biol (2022) doi:10.1016/J.SEMCDB.2022.05.012.

24. Farhy-Tselnicker, I. et al. Activity-dependent modulation of synapse-regulating genes in astrocytes. Elife 10, (2021).

25. Wilton, D. K., Dissing-Olesen, L. & Stevens, B. Neuron-Glia Signaling in Synapse Elimination. Annu Rev Neurosci 42, 107–127 (2019).

26. Chung, W. S., Allen, N. J. & Eroglu, C. Astrocytes Control Synapse Formation, Function, and Elimination. Cold Spring Harb Perspect Biol 7, (2015).

27. Bonetto, G., Belin, D. & Káradóttir, R. T. Myelin: A gatekeeper of activity-dependent circuit plasticity? Science 374, (2021).

28. Sancho, L., Contreras, M. & Allen, N. J. Glia as sculptors of synaptic plasticity. Neurosci Res 167, 17–29 (2021).

29. Allen, N. J. & Eroglu, C. Cell Biology of Astrocyte-Synapse Interactions. Neuron 96, 697– 708 (2017).

30. Baldwin, K. T. & Eroglu, C. Molecular mechanisms of astrocyte-induced synaptogenesis. Curr Opin Neurobiol 45, 113–120 (2017).

31. Tan, C. X., Burrus Lane, C. J. & Eroglu, C. Role of astrocytes in synapse formation and maturation. Curr Top Dev Biol 142, 371–407 (2021).

32. Watanabe, S. et al. Clathrin regenerates synaptic vesicles from endosomes. Nature 2014 515:7526 515, 228–233 (2014).

33. Begemann, I. & Galic, M. Correlative light electron microscopy: Connecting synaptic structure and function. Frontiers in Synaptic Neuroscience vol. 8 Preprint at 10.3389/fnsyn.2016.00028 (2016).

34. Staffler, B. et al. SynEM, automated synapse detection for connectomics. Elife 6, (2017).

35. Dorkenwald, S. et al. Automated synaptic connectivity inference for volume electron microscopy. Nature Methods 2017 14:4 14, 435–442 (2017).

36. Chandrasekaran, S. et al. Unbiased, High-Throughput Electron Microscopy Analysis of Experience-Dependent Synaptic Changes in the Neocortex. Journal of Neuroscience 35, 16450–16462 (2015).

37. Segev, A., Garcia-Oscos, F. & Kourrich, S. Whole-cell Patch-clamp Recordings in Brain Slices. J Vis Exp 2016, 54024 (2016).

38. Ippolito, D. M. & Eroglu, C. Quantifying Synapses: an Immunocytochemistry-based Assay to Quantify Synapse Number. JoVE (Journal of Visualized Experiments) e2270 (2010) doi:10.3791/2270.

39. Maidorn, M., Rizzoli, S. O. & Opazo, F. Tools and limitations to study the molecular composition of synapses by fluorescence microscopy. Biochem J 473, 3385–3399 (2016).

40. Blanco-Suarez, E., Liu, T. F., Kopelevich, A. & Allen, N. J. Astrocyte-Secreted Chordin-like 1 Drives Synapse Maturation and Limits Plasticity by Increasing Synaptic GluA2 AMPA Receptors. Neuron 100, 1116–1132.e13 (2018).

41. Stogsdill, J. A. et al. Astrocytic neuroligins control astrocyte morphogenesis and synaptogenesis. Nature 551, 192–197 (2017).

42. Holt, L. M. et al. Astrocyte morphogenesis is dependent on BDNF signaling via astrocytic TrkB.T1. Elife 8, (2019).

43. Nguyen, A. Q. et al. Astrocytic Ephrin-B1 Controls Excitatory-Inhibitory Balance in Developing Hippocampus. J Neurosci 40, 6854–6871 (2020).

44. Glebov, O. O., Cox, S., Humphreys, L. & Burrone, J. Neuronal activity controls transsynaptic geometry. Scientific Reports 2016 6:1 6, 1–11 (2016).

45. Baldwin, K. T. et al. HepaCAM controls astrocyte self-organization and coupling. Neuron (2021) doi:10.1016/j.neuron.2021.05.025.

46. Nguyen, A. Q. et al. Astrocytic Ephrin-B1 Controls Excitatory-Inhibitory Balance in Developing Hippocampus. J Neurosci 40, 6854–6871 (2020).

47. Verstraelen, P., Garcia-Diaz Barriga, G., Larsen, P. H., Timmermans, J.-P. & de Vos, W. H. Systematic Quantification of Synapses in Primary Neuronal Culture. iScience 23, 101542 (2020).

48. McNamara, G., Difilippantonio, M., Ried, T. & Bieber, F. R. Microscopy and Image Analysis. Curr Protoc Hum Genet 94, 4.4.1–4.4.89 (2017).

49. Dani, A., Huang, B., Bergan, J., Dulac, C. & Zhuang, X. Superresolution Imaging of Chemical Synapses in the Brain. Neuron 68, 843–856 (2010).

50. Risher, W. C. et al. Astrocytes refine cortical connectivity at dendritic spines. Elife 3, (2014).

51. Risher, W. C. et al. Thrombospondin receptor α2δ-1 promotes synaptogenesis and spinogenesis via postsynaptic Rac1. Journal of Cell Biology 217, 3747–3765 (2018).

52. Danielson, E., Lee, S. H. & Fox, M. A. SynPAnal: Software for rapid quantification of the density and intensity of protein puncta from fluorescence microscopy images of neurons. PLoS One 9, (2014).

53. Schmitz, S. K. et al. Automated analysis of neuronal morphology, synapse number and synaptic recruitment. J Neurosci Methods 195, 185–193 (2011).

54. Moreno Manrique, J. F., et al. SynapseJ: An Automated, Synapse Identification Macro for ImageJ. Front Neural Circuits 15, (2021).

55. Dzyubenko, E., Rozenberg, A., Hermann, D. M. & Faissner, A. Colocalization of synapse marker proteins evaluated by STED-microscopy reveals patterns of neuronal synapse distribution in vitro. J Neurosci Methods 273, 149–159 (2016).

56. Wang, Y. et al. SynQuant: an automatic tool to quantify synapses from microscopy images. Bioinformatics 36, 1599–1606 (2020).

57. Risher, W. C. et al. Thrombospondin receptor α2δ-1 promotes synaptogenesis and spinogenesis via postsynaptic Rac1. J Cell Biol 217, 3747 LP – 3765 (2018).

58. Irala, D. et al. Astrocyte-secreted neurocan controls inhibitory synapse formation and function. Neuron (2024) doi:10.1016/j.neuron.2024.03.007.

59. Berg, S. et al. ilastik: interactive machine learning for (bio)image analysis. Nat Methods 16, 1226–1232 (2019).

60. Eisele, R. Calculate the intersection area of two circles. https://www.xarg.org/2016/07/calculate-the-intersection-area-of-two-circles/ (2016).

61. Irala, D. et al. Astrocyte-Secreted Neurocan Controls Inhibitory Synapse Formation and Function 2. bioRxiv (2023) doi:10.1101/2023.04.03.535448.

62. Singh, S. K. et al. Astrocytes Assemble Thalamocortical Synapses by Bridging NRX1α and NL1 via Hevin. Cell 164, 183–196 (2016).

63. Allen, N. J. et al. Astrocyte glypicans 4 and 6 promote formation of excitatory synapses via GluA1 AMPA receptors. Nature 486, 410–414 (2012).

64. Christopherson, K. S. et al. Thrombospondins are astrocyte-secreted proteins that promote CNS synaptogenesis. Cell 120, 421–433 (2005).

65. Kucukdereli, H. et al. Control of excitatory CNS astrocyte-secreted protein. 43–45 (2016) doi:10.1073/pnas.1104977108.

66. Diniz, L. P. et al. Astrocyte-induced synaptogenesis is mediated by transforming growth factor β signaling through modulation of d-serine levels in cerebral cortex neurons. Journal of Biological Chemistry 287, 41432–41445 (2012).

67. Eroglu, Ç. et al. Gabapentin Receptor α2δ-1 Is a Neuronal Thrombospondin Receptor Responsible for Excitatory CNS Synaptogenesis. Cell 139, 380–392 (2009).

