## Supplemental Information for "SynBot: An open-source image analysis software for automated quantification of synapses"

**Table S1 SynBot Functions Defined Related to Figure 2**

| Step | Function Name | Description |
|---|---|---|
| 1a | 2-Channel Colocalization | The red and green image channels are analyzed for colocalizations. |
| 1b | 3-Channel Colocalization | The red, green, and blue image channels are analyzed for triple colocalizations. |
| 1- | Pick Channels | This option allows the user to change the color of the channels in the input images. This is particularly useful since SynBot requires the input images to have the channels to be analyzed in red, green, and blue. SynBot will attempt RGB conversion without this option selected and may have unintended results such as splitting magenta signal across red and blue or analyzing the wrong channels. |
| 2- | Noise Reduction (Recommended) | This option removes background by applying the FIJI Subtract Background function (rolling ball radius = 50) followed by a Gaussian Blur Filter (sigma = 0.57) to aid in object detection. |
| 2- | Brightness Adjustment | This option adjusts the image brightness using the FIJI Enhance Contrast function to have a certain percentage of saturated pixels. This option is not recommended unless images are consistently dim across conditions. Consider adjusting imaging parameters. |
| 3a | Manual | This thresholding method allows the user to manually threshold each channel of each image. |
| 3b | Fixed Value | This thresholding method allows the user to set a single threshold value for each channel that is used for all of the images being analyzed. |
| 3c | Percent Histogram | This thresholding method adjusts the image threshold such that the brightest pixels are kept. |

| | | |
|---|---|---|
| | | The percentage of pixels above threshold for each channel is provided by the user. |
| 3d | FIJI auto | This thresholding method allows the user to apply any of the automated thresholding algorithms in the FIJI auto threshold function. |
| 3e | ilastik | This thresholding method applies a user-trained random forest machine learning model for each channel to threshold the images. Training the model in ilastik is described in Berg et. al., 2019 *Nature Methods*. We recommend training the ilastik model on at least 5 images across conditions. |
| 3f | Pre-Thresholded | This thresholding method allows users to input pre-thresholded images into SynBot. |
| 3g | Threshold from File | This thresholding method allows users to enter a csv with threshold values to use for each channel of each image. Importantly, this feature can be used to replicate previous analyses and optimize other analysis parameters while keeping the thresholds the same. |
| 4a | Whole Image | This ROI option analyzes the whole input image. |
| 4b | Auto Cell Body | This ROI option can be used to identify a single cell body automatically and draw a circular ROI around it. This method does not work well if the image includes more than one cell. If this method fails SynBot will prompt the user to use the circle method. |
| 4c | Circle | This ROI option allows the user to select a central point for each image and creates a circle ROI around that point with a set radius. |
| 4d | Cell Territory | This ROI option can be used to automatically identify a cell’s territory using a cell fill marker in the blue channel. A mask is created which serves as the ROI for red and green colocalization. This |

| | | |
|---|---|---|
| | | ROI method only works with the 2-Channel colocalization method selected in step 1. |
| 4e | Custom | This ROI allows the user to set a custom ROI for each image. |
| 5 & 6 | Red/Green/Blue<br>Min/Max Pixel Size | This field allows the user to set a minimum and maximum pixel area to be counted as a punctum for each channel. |
| 7a | Circular-approximation | This analysis option calculates colocalizations by approximating red and green puncta to circles based on their radii and coordinates and using this to determine if they overlap. |
| 7b | Pixel-overlap | This analysis option uses the FIJI Image Calculator function to identify pixels that contain red and green signal that is above threshold and counts these as colocalizations. |
| 7- | 90-degree rotation control | This option rotates the red image 90 degrees to the right and then performs the SynBot analysis as usual. This can be used as a control to test if the number of colocalization observed are greater than what would be seen by chance given the abundance of the red and green markers. |
| | Experiment Directory | This opens a file explorer dialog to select the image input folder. |

**Table S2 Circularity of Synaptic Markers Related to Figures 5 and 6**

| Corresponding Figure | Marker | Average Circularity (Range 0-1) +/- standard deviation |
|---|---|---|
| 5 E & F | Homer1 | 0.882 +/- 0.0413 |

| | | |
|---|---|---|
| 5 E & F | Bassoon | 0.897 +/- 0.0162 |
| 5 G & H | Gephyrin | 0.884 +/- 0.0223 |
| 5 G & H | Bassoon | 0.911 +/- 0.0124 |
| 6 B & C | PSD95 | 0.922 +/- 0.0282 |
| 6 B & C | VGluT1 | 0.823 +/- 0.0314 |

**Figure S1 Raw Puncta Counts by Analysis Method Related to Figures 5 and 6**

**A)** Raw postsynaptic (red) puncta, presynaptic (green) puncta, and colocalized puncta counts for *In Vitro* Excitatory and *In Vitro* Inhibitory experiments related to figures 5E and 5G respectively. Lines indicate puncta counts for the same image across analysis methods. Control images are on the left of each plot marked with solid lines and ACM images are on the right side of each plot marked with dashed lines.

**B)** Raw postsynaptic (red) puncta, presynaptic (green) puncta, and colocalized puncta counts for *In Vivo* a2d1 experiments related to Figure 6C. Lines indicate puncta counts for the same image across analysis methods. WT images are on the left of each plot marked with solid lines and KO images are on the right side of each plot marked with dashed lines.

**C)** First Column: Two-way ANOVA analysis of data obtained by each experiment shown in figures 4, 5, and 6 testing for the effect of thresholding method on colocalized puncta count. Accompanying columns show pair-wise Tukey’s HSD post-hoc tests if the ANOVA was significant.

**Figure S2 Representative Images of *In Vitro* Synapses Counted by Each Thresholding Method Related to Figure 5**

- **A)** Separated Homer1 and Bassoon channels from zoom in excitatory synapse images shown in figure 5E. Scale: 15 µm for full image and 5 µm for zoom in.
- **B)** Zoom in images from figure 5E showing synapses detected by each thresholding method. Small white dots represent the synapses that were detected.
- **C)** Separated Gephyrin and Bassoon channels from zoom in inhibitory synapse images shown in figure 5G. Scale: 15 µm for full image and 5 µm for zoom in.
- **D)** Zoom in images from figure 5G showing synapses detected by each thresholding method. Small white dots represent the synapses that were detected.

**Figure S3 Colocalized Puncta Detected Using Circular Approximation vs. Pixel-by-pixel Analysis Types Related to Figure 6**

- A) Quantification of VGluT1-PSD95 colocalization using the SynBot manual thresholding method with either the circular approximation or pixel-by-pixel analysis types. Colocalized puncta counts were normalized to the WT average for each experimental pair. Mouse averages are shown as large black dots (N = 3 mice per condition) with individual images shown as small gray dots. Error bars represent 1 standard error of the mean.

A)

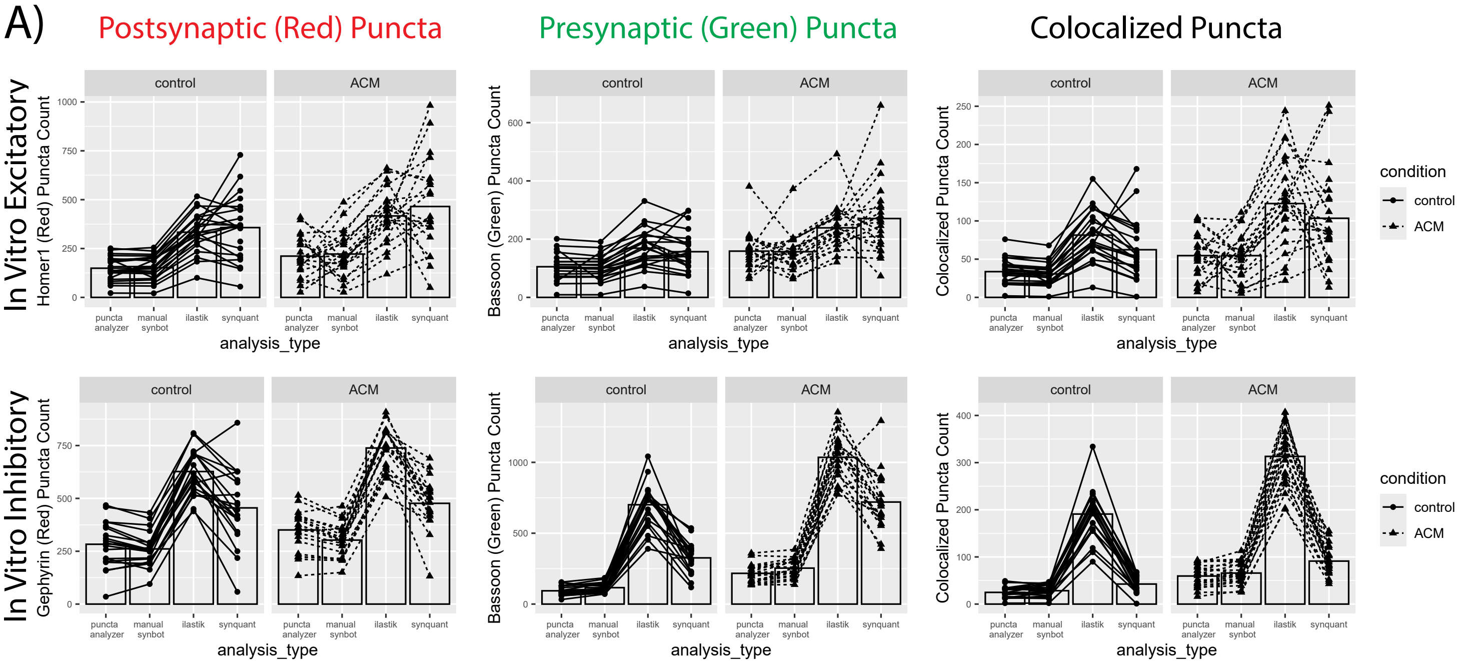


B)

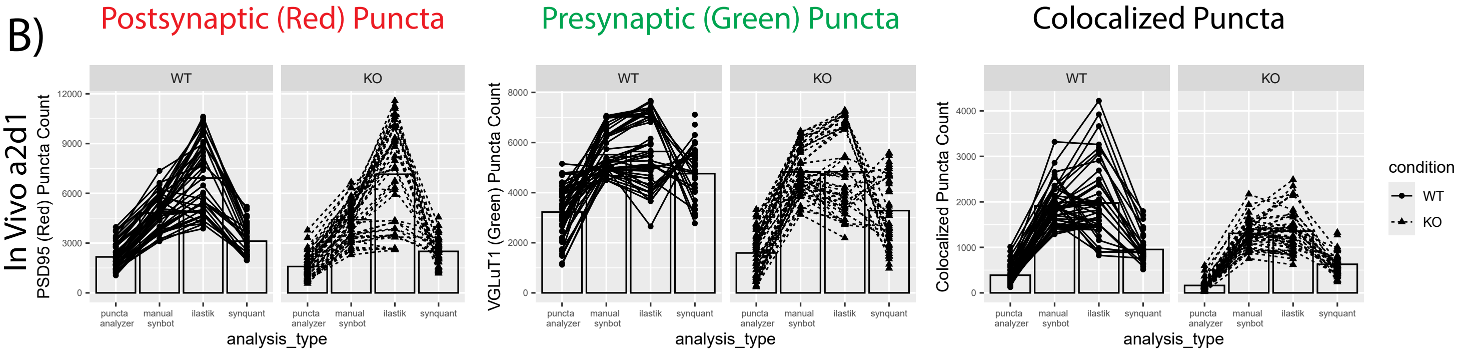


C)

| dataset | anova p value | manual_synbot-puncta_analyzer | ilastik-puncta_analyzer | synquant-puncta_analyzer | ilastik-manual_synbot | synquant-manual_synbot | synquant-ilastik |
|---|---|---|---|---|---|---|---|
| simulated_data_raw | 0.916 | | | | | | |
| invitro_excitatory_normalized | 0.896 | | | | | | |
| invitro_excitatory_raw | 4.87E-11 | 0.998 | 0 | 0 | 0 | 0 | 0.151 |
| invitro_inhibitory_normalized | 0.138 | | | | | | |
| invitro_inhibitory_raw | 2.00E-16 | 0.977 | 0 | 0.114 | 0 | 0.249 | 0 |
| invivo_excitatory_normalized | 0.05 | 0.096 | 0.083 | 0.135 | 0.999 | 0.999 | 0.997 |
| invivo_excitatory_raw | 2.00E-16 | 0 | 0 | 0 | 0.846 | 0 | 0 |

Figure S2

Savage et. al., 2024

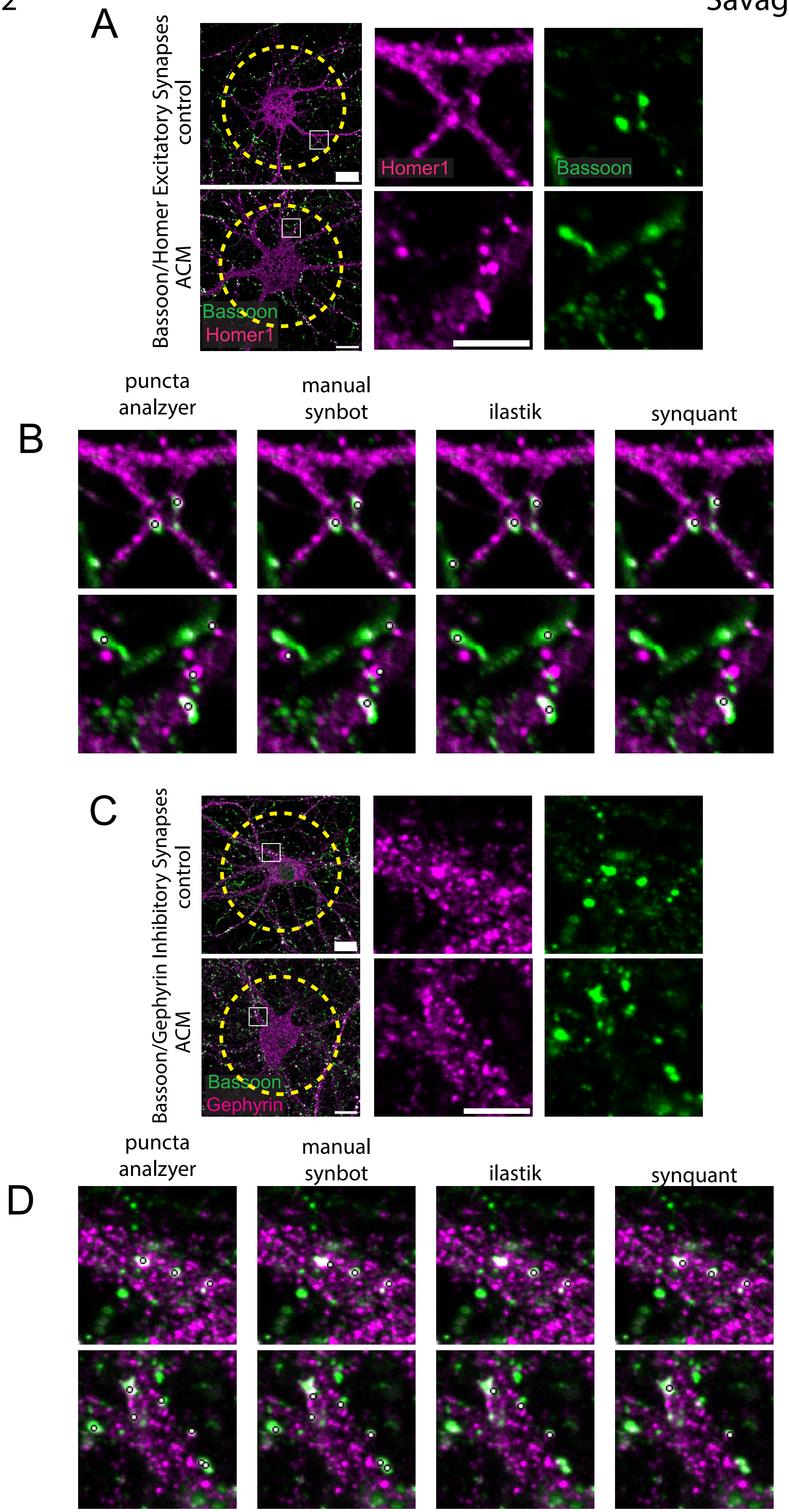

Figure S3

Savage et. al., 2024

A)

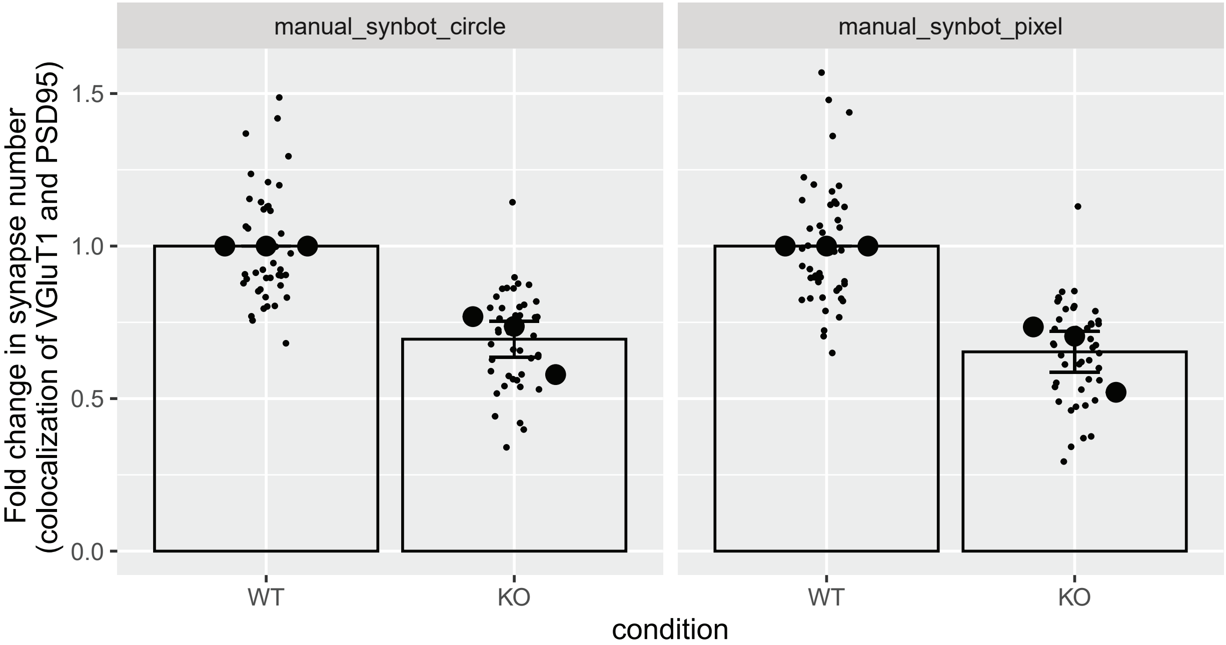
