## Supplementary material for "SynBot: An open-source image analysis software for automated quantification of synapses": Key Resources

**Key resources table**

| REAGENT or RESOURCE | SOURCE | IDENTIFIER |
|---|---|---|
| Antibodies | | |
| AffiniPure Goat anti-Rat IgG + IgM (H+L) | Jackson Immunoresearch | Cat# 112005044<br>RRID: AB_2338094 |
| AffiniPure Goat anti-Mouse IgG + IgM (H+L) | Jackson Immunoresearch | Cat# 115005044<br>RRID: AB_2338451 |
| Mouse anti-neural cell adhesion molecule L1 Hybridoma | Developmental Studies Hybridoma Bank | Cat# ASCS4<br>RRID: AB_528349 |
| Anti-Bassoon antibody | Enzo/Assay Designs | Cat# SAP7F07/VAM-PS003F<br>RRID: AB_2038857 |
| Anti-Gephyrin antibody | Synaptic Systems | Cat# 147002<br>RRID: AB_2619838) |
| Anti-Homer1 antibody | Synaptic Systems | Cat# 160002<br>RRID: AB_2120990 |
| Anti-VGAT antibody | Synaptic Systems | Cat# 131004<br>RRID: AB_887873 |
| Anti-Vglut1 antibody | Millipore | Cat# AB5905<br>RRID: AB_2301751 |
| Alexa Fluor 488 goat anti-Mouse IgG (H+L) | Invitrogen | Cat # A11001<br>RRID: AB_2534069 |
| Alexa Fluor 568 goat anti-Rabbit IgG (H+L) | Invitrogen | Cat# A11011<br>RRID: AB_143157 |
| Alexa Fluor 647 goat anti-Guinea pig IgG (H+L) | Invitrogen | Cat# A21450<br>RRID: AB_2535867 |
| Guinea pig anti- VGAT | Synaptic Systems | Cat# 131004<br>RRID: AB_887873 |
| Rabbit anti-Gephyrin | Synaptic Systems | Cat# 147002<br>RRID: AB_2619838 |
| Rabbit anti-PSD95 | Life Technologies | Cat# 51-6900<br>RRID: AB_2533914 |
| Chemicals, peptides, and recombinant proteins | | |
| 2,2,2-tribromoethanol | Sigma | Cat# T48402-25G |
| 2-methyl-2-butanol | Sigma | Cat# 152463-250mL |
| B27 | GIBCO | Cat# 17504044 |
| B27 Plus | GIBCO | Cat# A3582801 |
| BDNF | PeproTech | Cat# 450-02 |
| Boric Acid | Sigma | Cat# B0394 |
| BSA | Sigma | Cat# A4161 |
| BSL1 (Baneiraea Simplicifolia Lectin 1) | Vector Laboratories | Cat# L-1100 |
| CNTF | PeproTech | Cat# 450-13 |
| Cytosine arabinoside (AraC) | Sigma | Cat# C1768 |
| DAPI | Invitrogen | Cat#D1306 |
| DMEM | GIBCO | Cat# 11960 |
| DNaseI | Worthington | Cat# LS002007 |

| | | |
|---|---|---|
| DPBS with calcium, magnesium, glucose, and pyruvate | GIBCO | Cat# 14287 |
| DPBS without calcium or magnesium | GIBCO | Cat# 14190144 |
| Fetal Bovine Serum | Thermo Fisher | Cat# 10-437-028 |
| Forskolin | Sigma | Cat# F6886 |
| Glycerol | Acros Organics | Cat# 15892-0010 |
| Hydrocortisone | Sigma | Cat# H-0888 |
| Insulin | Sigma | Cat# 11882 |
| L-Glutamine | GIBCO | Cat# 25030-081 |
| Low protein binding tubes | Eppendorf | Cat# 022431081 |
| Mouse Laminin | Cultrex | Cat# 3400-010-01 |
| N-acetyl cysteine | Sigma | Cat# A8199 |
| n-Propyl gallate | Sigma | Cat# P3130-100G |
| Neurobasal | GIBCO | Cat# 21103049 |
| Neurobasal minus phenol red | GIBCO | Cat# 12348017 |
| Neurobasal Plus | GIBCO | Cat# A3582901 |
| Normal Goat Serum (NGS) | Thermo Fisher | Cat# 01-6201 |
| Optimal Cutting Temperature solution (OCT) | Tissue Tek | Cat# 4583 |
| Papain | Worthington | Cat# LK003178 |
| Pen/Strep | GIBCO | Cat# 15140 |
| PFA 16% | Electron Microscopy Sciences | Cat# 15710 |
| Poly-D-Lysine | Sigma | Cat# P6407 |
| Sodium Pyruvate | GIBCO | Cat# 11360-070 |
| Tris Base | VWR | Cat# 101174-856 |
| Triton X-100 | Roche | Cat# 11332481001 |
| Trypsin Inhibitor | Worthington | Cat# LS003083 |
| Critical commercial assays | | |
| Pierce BCA protein assay kit | Thermo Fisher | Cat# 23225 |
| Deposited data | | |
| Synapse microscopy images | This paper | DOI: 10.5281/zenodo.12191805 |
| Experimental models: Cell lines | | |
| Rat primary cortical neurons | This paper | N/A |
| Rat primary cortical astrocytes | This paper | N/A |
| Experimental models: Organisms/strains | | |
| Rat: Sprague-Dawley | Charles River | 001 |

| | | |
|---|---|---|
| Software and algorithms | | |
| SynBot (version 1.1.1) | This paper | Eroglu-Lab/Syn_Bot: Syn_Bot synapse calculation macro for FIJI (github.com)<br><br>https://doi.org/10.5281/zenodo.12192447 |
| FIJI (version 2.14.0) | NIH | https://fiji.sc/<br>RRID:SCR_002285 |
| Ilastik (version 1.3.3) | Anna Kreshuk's lab (EMBL) | https://www.ilastik.org/<br>RRID:SCR_015246 |
| Puncta Analyzer (version 2.0) | Ippolito and Eroglu, 2010 | https://github.com/toddstavish/puncta-analyzer |
| R: A Language and Environment for Statistical Computing (version 4.3.3) | R Core Team | https://cran.r-project.org/<br>RRID:SCR_001905 |
| R package nlme (version 3.1-164) | R Core Team | https://www.rdocumentation.org/packages/nlme/versions/3.1-162<br>RRID:SCR_015655 |
| Other | | |
| 20µm nylon mesh | Elko filtering | Cat# 03-20/14 |
| Vivaspin MWCO 5000; 20 mL tubes | Sartorius | Cat # VS2012 |
